# Aging restricts maturation of CXCL13^+^ T follicular helper cells in human immunity

**DOI:** 10.64898/2026.04.03.715992

**Authors:** Nathan A. Bracey, Casey Beppler, Tatjana Bilich, Adrienne H. Long, Elsa Sola, Aviv Barak, Timothy J. Few-Cooper, Azam Mohsin, Vishnu Shankar, Vamsee Mallajosyula, Lilit Kamalyan, Robson Capasso, Shai S. Shen-Orr, Mark M. Davis

**Affiliations:** Department of Medicine; Department of Microbiology, Immunology and Infectious Diseases, University of Calgary, Calgary, AB, Canada; Institute for Immunity, Transplantation and Infection, Stanford University School of Medicine, Stanford, CA, USA; Department of Pediatrics, Division of Hematology and Oncology, Stanford University, Stanford, CA, USA; Department of Immunology, Faculty of Medicine, Technion – Israel Institute of Technology, Haifa, Israel; Division of Sleep Surgery, Department of Otolaryngology-Head and Neck Surgery, Stanford University School of Medicine, Stanford, CA, USA; Department of Microbiology and Immunology, Stanford University School of Medicine, Stanford, CA, USA

**Keywords:** aging, Tfh cells, maturation, CXCL13, gene regulation, human immunology

## Abstract

A decline in specific antibody responses is a hallmark of human aging, yet the differential contributions of B and T lymphocytes and their interactions remain unclear. CXCL13 is a critical chemokine that shapes germinal center organization, but the regulation of human-specific CXCL13^+^ Tfh cells during aging is not known. Using human tonsil organoids, single-cell RNA sequencing, and CRISPR perturbations, we mapped age-associated changes in T follicular helper (Tfh) cells, the cell type that provides T cell “help” to B cells in germinal centers (GCs). Tonsil organoids from older donors generated weaker influenza-specific antibody responses, which we traced to Tfh cell defects rather than B cells. Single-cell profiling revealed a selective loss of mature CXCL13⁺ GC-Tfh cells accompanied by accumulation of Tfh precursor states. Trajectory analysis showed that aging arrests Tfh cell maturation at the early activated precursor transition, and CRISPR perturbations identified BACH2 and SOX4 as transcriptional regulators of differentiation reduced with age. These findings reveal a human-specific mechanism of immune aging with implications for strategies to restore humoral immunity.

## Introduction

The goal of vaccination is to elicit durable protective immunity through the generation of high-affinity antibodies. Although antibodies are produced by B cells, their quality depends on help from specialized CD4⁺ T cells known as T follicular helper cells. These Tfh cells interact with B cells in germinal centers of peripheral lymphoid organs, such as lymph nodes, tonsils, and spleen. Within these microenvironments, B cells proliferate and differentiate into antibody-secreting plasmablasts (PBs) and memory cells that together provide lasting immunity.^1–3^ Despite the strengths of vaccination, not all individuals are equally protected, and there is marked inter-individual variation in antibody responses. One of the most critical determinants of poor vaccine responsiveness is aging. Older adults generate fewer antibody-secreting B cells and produce reduced quantities of vaccine-specific antibodies compared to younger individuals, reflecting defects in immune responses that are not well understood.^4^

Previous work, stemming largely from mouse models, has shown that aging negatively impacts multiple aspects of GC biology that compromise the generation of durable antibody responses. GC-B cells from older mice show impaired class-switch recombination and somatic hypermutation, reduced activation-induced cytidine deaminase-driven programs, and diminished selection into long-lived plasma cell and memory pools compared to younger mice.^5^ In parallel, the quality of T cell help to B cells also declines with age. This has been variably attributed to mislocalization of Tfh cells within GCs,^6^ accumulation of precursor Tfh states without a corresponding increase in functional GC-Tfh cells,^7^ impaired late stage Tfh development,^8^ and unfavorable ratios of inhibitory T follicular regulatory (Tfr) cells to Tfh populations.^9^ In humans, aging has been similarly associated with weaker vaccine-specific antibody responses and fewer antibody-secreting cells after immunization,^4,10–13^ but the direct impact of aging on Tfh cells has been less clear. Analyses of circulating and lymphoid Tfh cell subsets have yielded conflicting results, with some studies reporting increased frequencies in older adults and others decreased.^7,9,14,15^ Notably, the human immune aging metric IMM-AGE, identified by longitudinal tracking of immune cell subset composition at high-resolution in peripheral blood, identified a reduction in circulating CD4⁺ Tfh cells as a key feature in the immune aging trajectory.^16^ Tfh cell biology is not identical across species. For example, some human GC-Tfh cells uniquely express the chemoattractant CXCL13, which is closely associated with GC activity and antibody quality.^17–19^ However, these cells are absent in mice, where CXCL13 is produced exclusively by follicular dendritic cells.^20^ Collectively, these findings highlight the B-Tfh axis as a critical point of vulnerability during aging, but the relative contributions of B and Tfh cells to impaired human antibody responses remain unresolved.

Tonsil-derived immune organoids provide a unique platform to explore human-specific immune function. They enable investigation of the cellular and molecular mechanisms that govern B and T cell interactions directly in human lymphoid tissue. Immune organoids recapitulate key features of GC responses, including B-Tfh cell interactions, plasmablast differentiation, somatic hypermutation of antibody genes, affinity maturation, and class switching, while preserving the immunological diversity of human lymphoid tissue.^21^ This system has already yielded insights into early immune development,^22^ the roles of principal regulatory T cell types,^23^ and vaccine-induced responses to influenza antigens,^24^ underscoring its utility for identifying mechanisms of adaptive immunity. Furthermore, immune organoids allow us to test hypotheses and define mechanisms independently of mouse models, which is critical to understanding human specific phenomena. Here, we performed an in-depth analysis of how aging shapes human immune responses using tonsil samples from donors aged 5-73 years. Organoids from older individuals mounted weaker antibody responses, which we traced to Tfh cell defects. Single-cell RNA sequencing (scRNA-seq) revealed a selective reduction of CXCL13⁺ GC-Tfh effector cells. Trajectory analysis and arrayed CRISPR-based perturbations demonstrated that aging arrests Tfh maturation and identified critical transcriptional regulators that are impaired in older donors. Together, these findings reveal a human-specific mechanism by which aging impacts the generation of high-quality antibody responses and uncover regulatory nodes that may be leveraged to restore humoral immunity in older adults.

## Results

### Humoral responses are impaired during human aging

The human immune system demonstrates substantial inter-individual variation, and one of the strongest predictors of adaptive immune responsiveness is age.^16,25–29^ To explore how aging shapes humoral immunity, we analyzed antibody responses to the seasonal influenza vaccine in a cohort of 402 healthy individuals aged 8-90 years.^24,30^ Peripheral blood was collected at baseline (day 0) and 28 days following vaccination, and neutralizing antibodies against hemagglutinin (HA) were quantified using the hemagglutination inhibition (HAI) assay.

In this cohort, vaccination elicited an increase in serum antibody titers, with most individuals showing measurable HAI response by day 28. Antibody titers to the H1, H3, and B vaccine components aggregated as a composite geometric mean titer (GMT) showed a 3-fold median change from baseline (**Figure 1a**). Aging emerged as a major determinant of vaccine responsiveness. Age negatively correlated with day 28 fold-change in GMT, corresponding to ∼4% decrease in HAI per decade of life (β=-0.006 per year, *P*<0.001). To further resolve how aging influences the quality of humoral immunity, we examined seroconversion (SC, defined as ≥4-fold rise and day 28 GMT ≥40) and seroprotection (SP, day 28 GMT ≥ 40) as complementary metrics capturing distinct aspects of vaccine immunogenicity. Previous human studies have suggested that pre-existing antibodies are an important determinant of subsequent vaccine responses, and while older adults maintain baseline antibody levels, they demonstrate reduced capacity to boost antibody titers following vaccination.^31,32^ Consistent with this, the probability of seroconversion declined with age (*P*<0.001, odds ratio (OR) per decade=0.86), whereas seroprotection was only modestly impacted (*P*=0.017, OR=0.88; **Figure 1b,c**). Collectively, these results support the notion that older individuals may retain pre-existing antibody-secreting cells but have limited ability to boost existing or generate de novo antibody responses. This distinction between antibody maintenance and induction suggests that aging does not uniformly impair all aspects of humoral immunity, but rather impacts the complex cellular interactions required for the generation and amplification of antibody responses.

**Figure 1.**
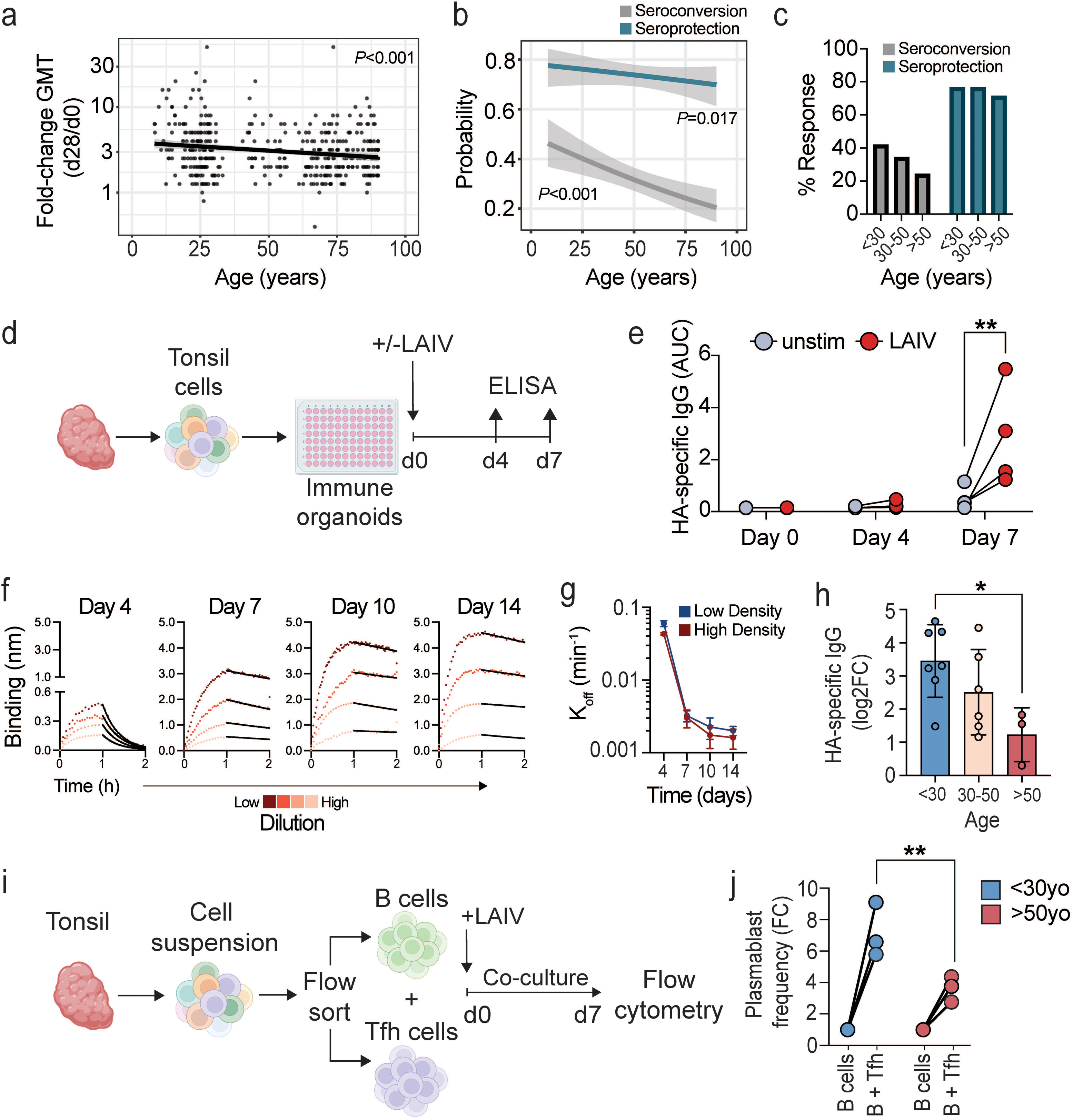
Human humoral immune responses decline with age. **a,** Linear regression analysis examining the relationship between the day 28 fold-change in geometric mean titer (GMT) using hemagglutination inhibition assay (y-axis) and age (x-axis). Each dot represents one donor (n = 402). **b,** Logistic regression analysis examining the probability of seroconversion (SC, defined as ≥ 4-fold rise and day 28 GMT ≥ 40) and seroprotection (SP, day 28 GMT ≥ 40) vs. age. **c,** Seroconversion (grey) and seroprotection (blue) in relation to donor age. **d,** Schematic representation of human tonsil immune organoids. Single-cell tonsil suspensions were embedded in a hydrogel matrix and cultured in 96-well plates for 7 days with LAIV. Supernatants were collected on day 0, 4, and 7. **e,** HA-specific IgG measured by ELISA from day 0, 4, and 7 organoid supernatants (n = 4) stimulated with LAIV (red) vs. unstimulated organoid supernatants (grey). Data points represent the area under the curve (AUC). Two-way ANOVA with Dunnett’s multiple comparisons test. **f,** Biolayer interferometry sensorgrams depicting the real-time binding kinetics of polyclonal HA-specific antibodies from tonsil cultures on day 4 through day 14 after stimulation with LAIV. The curves represent the binding response (nm shift) for different concentrations of HA-specific antibodies over time. Each measurement was performed in duplicates. **g,** Dissociation rate constant K_off_ per minute under low-density and high-density organoid conditions measured in duplicates. **h,** HA-specific IgG measured by ELISA from day 7 organoid supernatants in relation to donor age (n = 16). Data points represent the log2 fold change (FC) relative to matched day 7 unstimulated controls. One-way ANOVA with post-hoc Tukey test. **i,** Schematic representation of B-Tfh co-culture assay. B cells and Tfh cells were sorted from tonsil single-cell suspensions. Co-cultures were stimulated with LAIV for 7 days before measurement of plasmablast frequencies using flow cytometry. **j,** Plasmablast frequency of three younger (blue, left) and three older (red, right) donors shown as fold change (FC) relative to B cell only control. Data in **e** and **h** were analyzed using one-way ANOVA with post-hoc Tukey test. Data in **j** were analyzed using two-way ANOVA with Šídák’s multiple comparisons test. \**P* < 0.05, \*\**P* < 0.01.

To further explore the effects of aging on humoral immune responses directly within human peripheral lymphoid tissues, we used our recently described tonsil immune organoid model and adapted it to a 96-well plate format for improved throughput by embedding cells in hydrogel (**Figure 1d**).^21^ Stimulation of immune organoids with Live Attenuated Influenza Vaccine (LAIV) induced robust secretion of HA-specific IgG antibodies after 7 days (**Figure 1e**). Biolayer interferometry revealed increasing affinities of polyclonal HA-specific antibodies throughout 14 days of culture, confirming ongoing affinity maturation (**Figure 1f,g**).

We next assembled immune organoids from tonsil donors spanning almost 7 decades of life (n=16, 16-73 years, **Table S1**) and measured their responses to LAIV. Consistent with our *in vivo* human influenza vaccination studies, organoid immune responses were significantly reduced with age.^4,10–13^ Organoids generated from younger donors (<30 years) mounted robust responses to LAIV, with a strong induction of HA-specific IgG compared to unstimulated controls (**Figure 1h**). In contrast, organoids from older adults (>50 years) showed a significantly reduced vaccine response.

Previous work associated reduced humoral immune responses in aging with a decreased ratio of Tfh cells to inhibitory Tfr cells.^9^ Consistent with this, we found that tonsils from older adults contained increased frequencies of Tfr cells and reduced frequencies of GC-Tfh cells compared to younger donors (**Figure S1a,b**).^9^ Although these quantitative changes suggest an altered T cell landscape, their functional impact on GC activity in human tissues remains poorly defined. To test whether Tfr activity was required for the impaired B cell differentiation to antibody-secreting plasmablasts with age, we established Tfh-B cell co-cultures with normalized input ratios using sorted CXCR5^hi^PD-1^hi^ GC-Tfh cells and autologous tonsillar B cells from both young and old donors (**Figure 1i**). Cells were stimulated with LAIV and after 7 days we measured differentiation to CD38^hi^CD27^hi^ plasmablasts by flow cytometry (**Figure S1c**). In younger donors, the addition of Tfh cells to B cells elicited a strong helper response, with a 4-9-fold increase in plasmablast frequencies relative to B cells cultured alone (**Figure 1j**). Strikingly, even in the absence of Tfr cells, equalizing the Tfh cell input failed to completely restore the response in older donors. Plasmablast induction was markedly reduced, showing only a 3-fold increase. Together, these results demonstrate that the impact of aging on humoral immunity cannot be explained solely by shifts from Tfh to Tfr subsets. Rather, aging appears to attenuate the key role of Tfh cell help in enabling robust B cell responses.

### B cells maintain responsiveness to T cell help during human aging

Since immune organoids from older adults failed to mount strong antibody responses, and normalizing the frequencies of B and Tfh cells did not restore the defect, we next asked whether aging impairs B cell responsiveness to T cell-derived signals. To address this, we isolated B cells from peripheral blood mononuclear cells (PBMCs) of older and younger donors (n=20, 22-72 years). We cultured purified B cells with or without B cell receptor (BCR)-activating signals (IgA/IgG/IgM crosslinker) and/or signals mimicking T cell help (IL-21, CD40L) to assess age-related changes to B cells (**Figure 2a**). After 7 days, we measured the frequencies of differentiated B cell subsets using flow cytometry (**Figures 2b and S2**). Interestingly, B cells from both older and younger individuals responded similarly to exogenous BCR and T cell-derived signals. The frequencies of GC-like B cells and plasmablasts both showed similar increases compared to unstimulated conditions (**Figure 2c,d**). Similarly, CD138 upregulation and IgD/CD95 expression showed no significant differences between age groups under any conditions tested (**Figures 2e-h**). These results indicate that age-related decline in antibody induction is not primarily attributable to an intrinsic defect in B cell activation or differentiation. Instead, these data suggest that T cells have the dominant role in the attenuation of humoral immunity with age, consistent with murine studies.^33,34^

**Figure 2.**
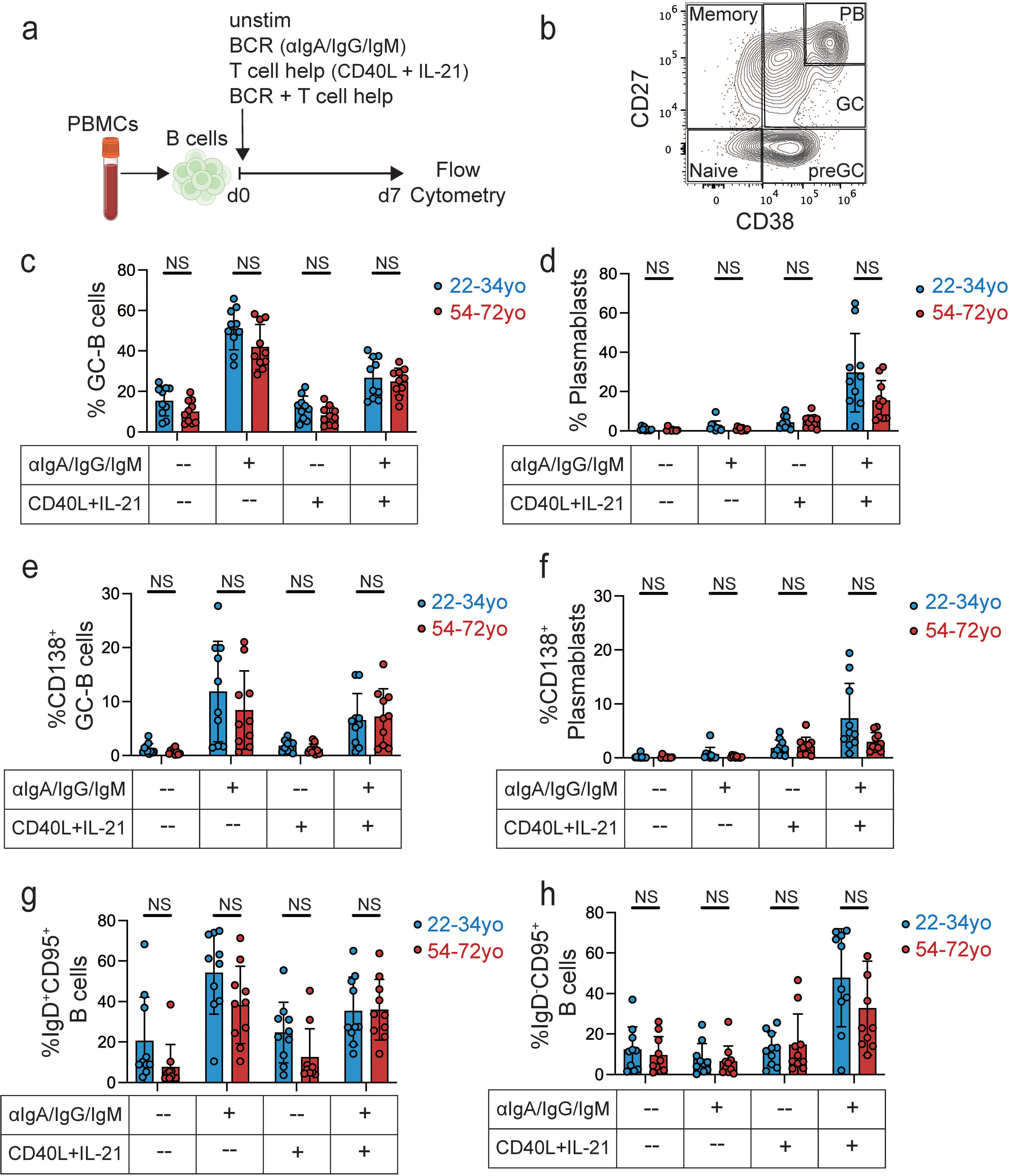
Responsiveness of B cells to T cell-derived signals is maintained during aging. **a,** Schematic representation of the protocol for measuring B cell responsiveness to *in vitro* B cell receptor (BCR) activation and T cell help signals. **b,** Exemplary gating for germinal center (GC)-B cells (CD27^mid^CD38^mid^) and plasmablasts (PB; CD27^hi^CD38^hi^) from CD19^+^, viable, single cells. **c-h,** Induction of different B cell phenotypes by different stimulus combinations using samples from 22-34yo (blue, n = 10) vs. 54-72yo donors (red, n = 10) for GC-B cells (**c**), plasmablasts (**d**), CD138^+^ GC-B cells (**e**), CD138^+^ plasmablasts (**f**), IgD^+^CD95^+^ B cells (**g**), and IgD^-^CD95^+^ B cells (**h**). Data were analyzed using Šídák’s multiple comparisons test (**c-h**). Stimulation was confirmed using two-way ANOVA (*P* < 0.0001). NS, not significant.

### CXCL13^+^ GC-Tfh cells are reduced during human aging

Since B cells from older adults responded normally to exogenous T cell-derived signals, we hypothesized that age-associated defects in humoral immunity might instead originate within the Tfh cells themselves. To investigate how aging influences the composition and molecular features of human tonsillar T cells, we purified CD3^+^ T cells from donors across a broad age range and performed scRNAseq (**Figures 3a and S3a**). After initial quality control, we identified 20,657 single-cell transcriptomes (**Figures S3b-d**), which clustered into 15 distinct populations representing canonical CD4^+^, CD8^+^, and unconventional T cell subsets (**Figures 3b and S3e**). CD4^+^ T cells were predominantly composed of naïve and central memory subsets, with smaller proportions of effector memory and activated cells. A distinct fraction exhibited hallmarks of Tfh cells, defined by elevated expression of *BCL6, CXCR5,* and *PDCD1*, consistent with localization and function within germinal centers.

**Figure 3.**
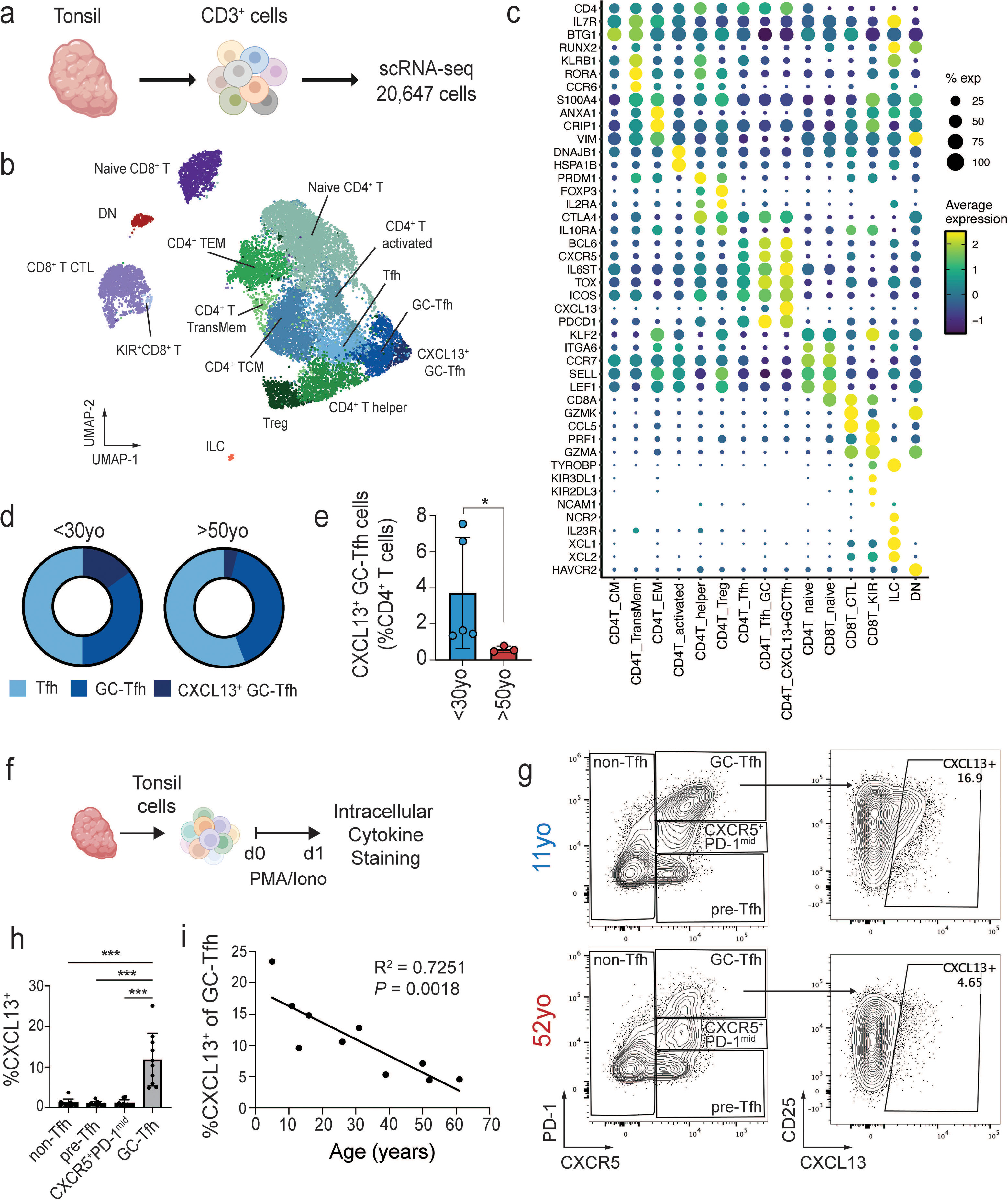
CXCL13^+^ GC-Tfh cells are reduced during human aging. **a,** Schematic workflow for processing human tonsil tissue to enrich T cells for single-cell RNA sequencing (scRNA-seq). **b,** Uniform Manifold Approximation and Projection (UMAP) of tonsil T cell transcriptomes. Each color represents one of 15 distinct T cell clusters. **c,** Average expression (dot color) and percent expression (dot size) are shown for gene markers used in cluster annotation. **d,** Overview of the proportional representation of Tfh (light blue), GC-Tfh (blue), and CXCL13^+^ GC-Tfh (dark blue) subsets among the three Tfh clusters in younger (left) vs. older donors (right). **e,** Frequencies of CXCL13^+^ GC-Tfh cells in tonsil samples of younger (blue, n = 5) and older (red, n = 3) donors. **f,** Schematized protocol for measuring Tfh cell subsets using flow cytometry and intracellular cytokine staining. **g,** Exemplary intracellular staining of CXCL13 in GC-Tfh cells (viable, CD45^+^CD19^-^CD14^-^CD16^-^CD8^-^CXCR5^+^PD-1^hi^) for one young (blue, top) and one old donor (red, bottom). **h,** Frequency of CXCL13^+^ cells in non-Tfh (CXCR5^-^), pre-Tfh (CXCR5^+^PD-1^lo^), CXCR5^+^PD-1^mid^ Tfh, and CXCR5^+^PD-1^hi^ GC-Tfh cell subsets by intracellular staining. Each dot represents one donor (n = 10). **i,** Linear regression analysis examining the correlation between the percentage of CXCL13^+^ GC-Tfh cells (y-axis) and age (x-axis). Each dot represents one donor (n = 10). Data in **e** were analyzed using Mann Whitney test. Data in **h** were analyzed using one-way ANOVA with Dunnett’s multiple comparisons test. \**P* = 0.0357, \*\*\**P* < 0.001.

Importantly, we identified three transcriptionally distinct populations of tonsil Tfh cells (**Figure 3c**), including early Tfh cells with lower *CXCR5, TOX*, and *BCL6* expression; classical GC-Tfh cells with high *BCL6, CXCR5*, and *PDCD1* expression; and a small but distinct subset of CXCL13^+^ GC-Tfh cells characterized by the highest expression of *BCL6* and *IL6ST* along with robust *CXCL13* transcription. To determine how these populations are affected by age, we compared the Tfh subset composition between younger and older donors, revealing a striking reduction of CXCL13^+^ GC-Tfh cells in older individuals (**Figure 3d,e**). In younger donors, CXCL13^+^ GC-Tfh cells comprised 1-8% of all CD4^+^ T cells, whereas in older individuals they accounted for <1%.

To validate the relationship between age and CXCL13^+^ GC-Tfh cells, we next analyzed an independent cohort of tonsil donors (n = 10, ages 5-61) by intracellular cytokine staining (ICS) (**Figure 3f,g**), using CXCR5 and PD-1 as previously established markers of Tfh cell maturation (**Figure S3f**).^7,35^ CXCL13 expression was restricted to the CXCR5^+^PD-1^hi^ GC-Tfh population and was absent from non-Tfh (CXCR5^-^), pre-Tfh (CXCR5^+^PD-1^lo^), and CXCR5^+^PD-1^mid^ Tfh subsets (**Figure 3h**). Notably, only a subset of GC-Tfh cells expressed CXCL13 (4-23%), suggesting that these cells may represent a distinct effector state within the GC-Tfh lineage. Consistent with our previous scRNA-seq data, the frequency of CXCL13^+^ GC-Tfh cells declined significantly with age (R^2^=0.73, *P*=0.0018, **Figures 3i and S3g**). CXCL13 secretion measured by ELISA showed a similar, though weaker, negative association with age (R^2^=0.34, *P*=0.0783), whereas secretion of the key Tfh effector cytokine IL-21 remained unchanged (R^2^=0.0998, *P*=0.3740), implying that its regulation is separate from that of the CXCL13^+^ GC-Tfh program (**Figure S3h**).

As a final validation using the publicly available human tonsil atlas dataset,^36^ we identified the population corresponding to *CXCL13* expression (annotated as GC-Tfh-SAP; **Figure S4a**) and confirmed its reduction with age (**Figure S4b**). Interestingly, *IL21* was again differentially regulated relative to *CXCL13*, with increased expression of *IL21* observed in a distinct Tfh population (annotated as GC-Tfh-OX40; **Figure S4c**).

Collectively, these data reveal that aging is associated with a loss of CXCL13^+^ GC-Tfh cells, distinguished by high *BCL6* and *IL6ST* expression and distinct effector potential, thus highlighting this uniquely human population as a likely key mediator of Tfh-dependent humoral immunity.

### CXCL13^+^ GC-Tfh cells constitute a distinct Tfh effector program

Tfh cell maturation is a multistep process that is shaped by interactions with both dendritic cells and B cells within developing GCs. Mouse models have provided a foundational framework for understanding Tfh cell development, however key differences exist between species. In mice, mature GC-Tfh cells upregulate CXCR5 and PD-1, but do not express CXCL13 (**Figure 4a**).^3^ In contrast, our data suggest that human Tfh cell maturation is more complex and involves an additional effector stage characterized by CXCL13 expression.

**Figure 4.**
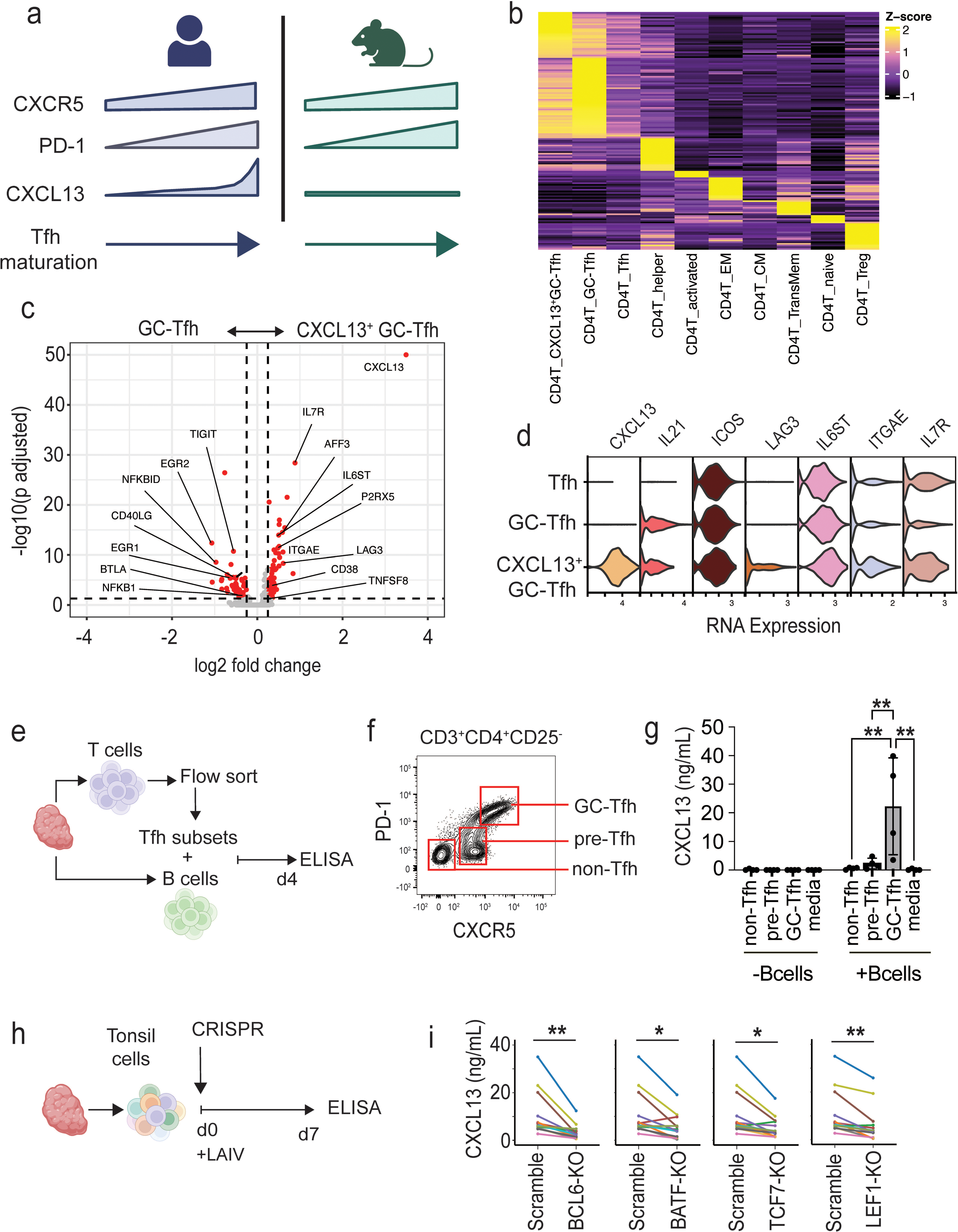
CXCL13^+^ GC-Tfh cells constitute a distinct Tfh effector program. **a,** Schematics of key differences in Tfh cell maturation between humans (left) and mice (right). **b,** Gene expression signatures of all CD4^+^ T cell clusters from tonsillar T cell scRNA-seq. **c,** Volcano plot showing differential gene expression of GC-Tfh cells (left) vs. CXCL13^+^ GC-Tfh cells (right). **d,** Select subset-specific marker gene expression in Tfh, GC-Tfh, and CXCL13^+^ GC-Tfh cells. **e,** Schematic overview for assessment of CXCL13 production by non-, pre-, and GC-Tfh cells. **f,** Exemplary gating for non-Tfh (PD-1^-^CXCR5^-^), pre-Tfh (PD-1^lo^CXCR5^lo^), and GC-Tfh cells (PD-1^hi^CXCR5^hi^). **g,** CXCL13 secretion by different sorted T cell subtypes cultured with or without B cells (n = 4). **h,** Schematic representation for CRISPR/Cas9-based editing of tonsil samples. **i,** CXCL13 ELISA of tonsillar cell culture supernatants for indicated transcription factor knockouts vs. scramble control (n = 14; ≤ 30 yo). Data were analyzed using two-way ANOVA with Šídák’s multiple comparison test (**g**) and one-way ANOVA with Dunnett’s multiple comparisons test (**i**). \**P* < 0.05, \*\**P* < 0.01.

To better define these CXCL13^+^ GC-Tfh cells, we analyzed our scRNA-seq tonsillar T cell dataset and compared gene expression profiles across all CD4^+^ T cell clusters. Interestingly, CXCL13^+^ GC-Tfh cells demonstrated a distinct transcriptomic signature compared to developing Tfh and GC-Tfh cells (**Figure 4b**). Differentially expressed genes upregulated in CXCL13^+^ GC-Tfh cells compared to other GC-Tfh cells included regulators of survival and metabolism (*GIMAP5, GIMAP7, GIMAP2, BCL2L11, SESN1, BCAT1*), markers of activation and GC residency (*ITGAE, LAG3, CD38*), T cell signaling molecules (*IL6ST, IL7R, FYB*), and transcriptional regulators (*AFF3, ARID5A, TCF7*; **Figure 4c**). In contrast, classical GC-Tfh cells were enriched for *TIGIT, EGR1, EGR2, CD40LG, BTLA4*, and *NFKB1*, reflecting a transcriptional bifurcation within the GC-Tfh compartment. This was further supported by progressive increases in key effector genes (*ICOS, IL6ST, ITGAE*), with CXCL13^+^ GC-Tfh cells showing the highest expression of these activation- and Tfh signaling-associated transcripts (**Figure 4d**). To determine whether the CXCL13^+^ GC-Tfh effector program corresponds to any established mouse Tfh subsets, we compared published murine Tfh cell gene signatures with CD4^+^ T cell transcripts from our scRNA-seq dataset.^37–41^ Mouse Tfh signatures generally exhibited minimal overlap with non-Tfh CD4^+^ T cells, and their enrichment was greatest in conventional GC-Tfh cells, suggesting that the CXCL13^+^ GC-Tfh subset represents a distinct, human-specific effector state (**Figure S5**).

We next examined the functional requirements for sustaining the CXCL13^+^ GC-Tfh effector program *in vitro*. We flow-sorted distinct Tfh subsets and cultured them in the presence or absence of autologous tonsillar B cells (**Figure 4e,f**). Consistent with our previous ICS experiments, only CXCR5^+^PD-1^hi^ GC-Tfh cells produced measurable CXCL13 (**Figure 4g**). In the absence of B cells however, CXCL13 production was completely abolished, indicating that maintenance of the CXCL13^+^ effector state requires continuous B-cell interactions.

To functionally validate that CXCL13 secretion is driven by core Tfh programming, we next performed targeted CRISPR knockout of canonical Tfh-associated transcription factors (TFs) in primary tonsil cells, followed by stimulation with LAIV for 7 days (**Figure 4h**). Disruption of *BCL6, BATF, TCF7,* and *LEF1* each significantly reduced CXCL13 secretion compared to scramble controls (**Figure 4i, Tables S2-3**), confirming their essential roles in sustaining the CXCL13^+^ GC-Tfh effector phenotype.

Together, these results establish CXCL13^+^ GC-Tfh cells as a late stage B cell-dependent effector population within the human Tfh lineage. This subset is transcriptionally distinct from other GC-Tfh cells and uniquely adapted to express the chemoattractant molecule CXCL13 within the germinal center during humoral immune responses.

### GC-Tfh cell maturation is impaired during human aging

Having established that CXCL13^+^ GC-Tfh cells represent a distinct effector state that is underrepresented in older donors, we next asked whether aging alters the developmental programming that gives rise to these cells. Flow cytometry profiling revealed that pre-Tfh cells (CXCR5^+^PD-1^lo^), which represent the earliest stage of Tfh maturation, increased significantly with age (R^2^=0.5476, *P*=0.0092; **Figures 5a and S6a,b**). This finding is in keeping with Webb et al., who showed that pre-Tfh cells are overrepresented in aged mouse lymph nodes following viral infection.^7^ In contrast, more advanced Tfh subsets declined with age (CXCR5^+^PD-1^mid^: R^2^=0.4653, *P*=0.0208; GC-Tfh: R^2^=0.5101, *P*=0.0135). Notably, ICOS expression was also negatively correlated with age (R^2^=0.5375, *P*=0.0103), consistent with the loss of CXCL13^+^ GC-Tfh cells and impaired expression of this CXCL13^+^ effector program gene that mediates T cell help to B cells (**Figure S6c**). These findings indicate that the accumulation of precursor states comes at the expense of mature GC-Tfh cell frequencies.

**Figure 5.**
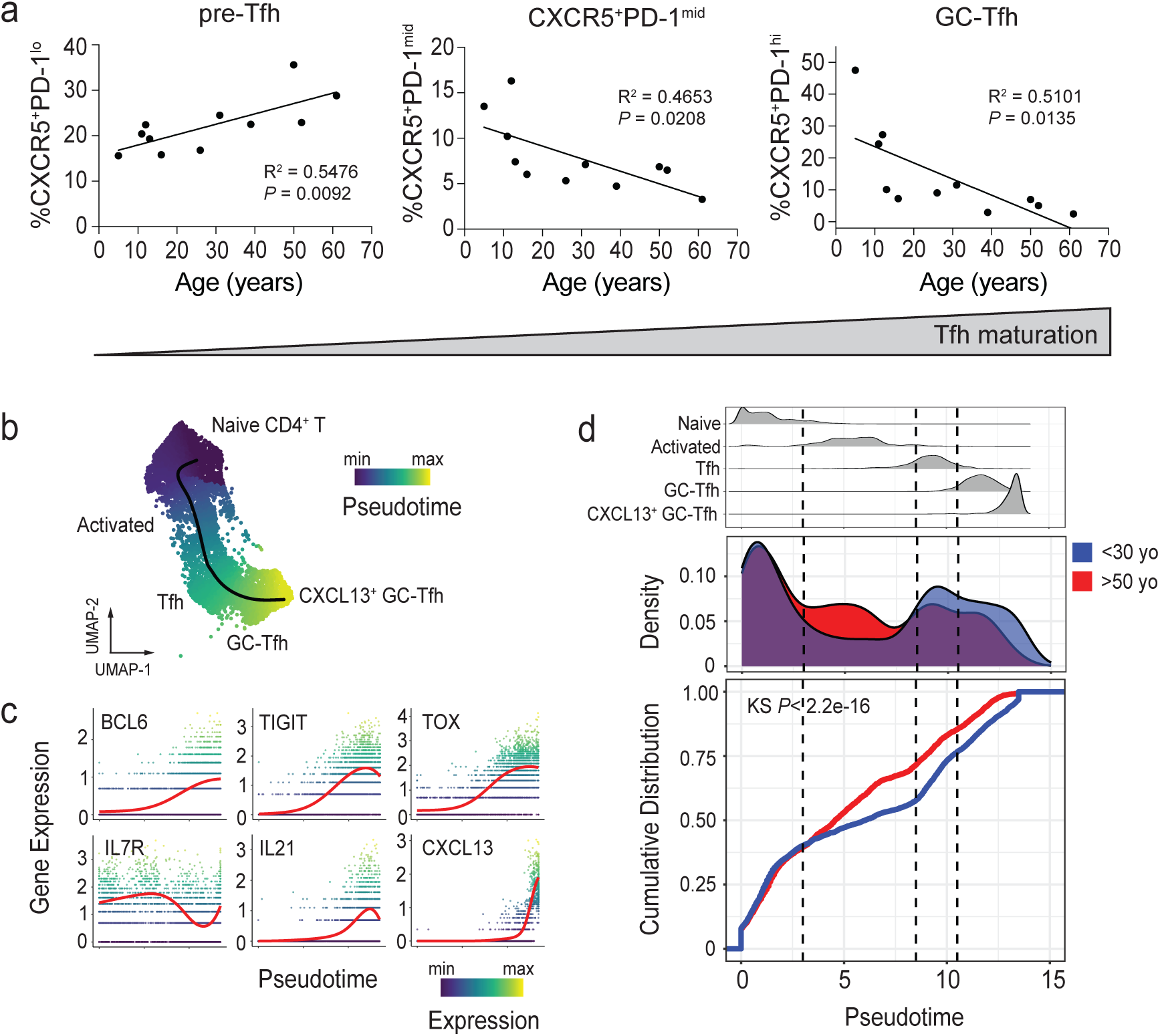
Aging is associated with impaired GC-Tfh cell maturation. **a,** Linear regression analysis examining the correlation of age (x-axis) and the percentage of Tfh cells within the CD4^+^ T cell compartment (y-axis) along their activation-to-differentiation trajectory from CXCR5^+^PD-1^lo^ (left), over CXCR5^+^PD-1^mid^ (middle), to mature GC-Tfh cells (right). Gated on viable CD3^+^CD19^-^CD4^+^FOXP3^-^. Each dot represents one donor (n = 11). **b,** UMAP representation of pseudotemporal trajectory from naïve CD4^+^ T cells (top left) to CXCL13^+^ GC-Tfh cells (bottom right) colored by pseudotime (blue to yellow color scale). **c,** Kinetics of select marker gene expression over pseudotime during Tfh cell maturation. **d,** Distribution of Tfh cell subsets (top) aligned with probability density (middle) and cumulative distribution (bottom) of Tfh cells across pseudotime in younger (blue) vs. older (red) donors.

To further explore this developmental pathway, we constructed a pseudotemporal trajectory from naïve CD4^+^ T cells to CXCL13^+^ GC-Tfh cells using our tonsillar scRNA-seq dataset (**Figure 5b**). Along this continuum, we observed a coordinated up-regulation of transcription factors (*BCL6, TOX*), co-inhibitory receptors (*TIGIT*), and Tfh effector molecules (*IL21, CXCL13*; **Figure 5c**). Comparing pseudotime distributions across age groups revealed that despite similar representations of naïve CD4^+^ T cells, older individuals accumulated cells at the activated CD4^+^ T cell state and were depleted at the mature GC-Tfh endpoints (**Figure 5d**). Analysis of the cumulative distribution (showing the fraction of cells that have reached a particular cell state across pseudotime) confirmed a divergence between younger and older donors emerging at the activated CD4^+^ T cell stage and persisting throughout later steps of maturation (Kolmogorov-Smirnov test, *P*<2.2e-16). Together, these data demonstrate that aging limits progression along the Tfh developmental pathway, producing a bottleneck in early T cell activation and pre-Tfh cell states.

### Knockout of age-dependent TFs reveals novel regulators of human Tfh cell maturation

To identify the molecular features underlying this impaired Tfh cell development during aging, we compared gene expression between younger and older individuals across pseudotime bins. Strikingly, most age-associated differences emerged at early differentiation states, even preceding GC-Tfh commitment (**Figure 6a**). This finding suggests that reduced frequencies of effector GC-Tfh cells observed in older individuals could result from changes that were already present within naïve CD4^+^ T cells recruited to lymphoid tissues.

**Figure 6.**
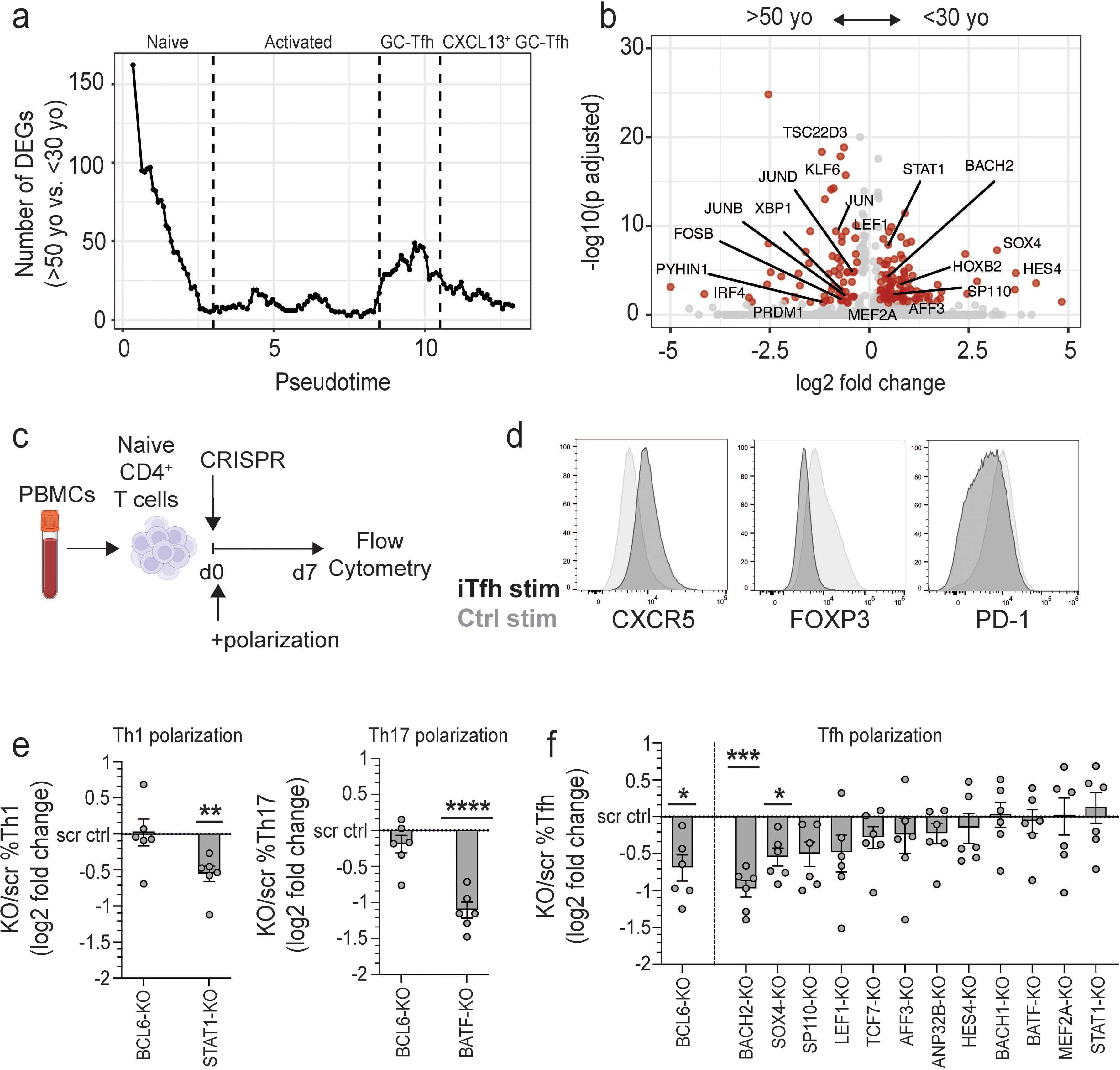
Knockout of age-dependent transcription factors reveals novel regulators of human Tfh cell differentiation. **a,** Number of differentially expressed genes (DEGs, >50yo vs. <30yo) over pseudotime during Tfh cell development from naïve CD4^+^ T cells (left) to CXCL13^+^ GC-Tfh cells (right). False discovery rate (FDR) is set to < 0.05. **b,** Volcano plot showing differential gene expression of Tfh cells from older (left) vs. younger donors (right). **c,** Schematic representation for CRISPR/Cas9-based editing of naïve CD4^+^ T cells followed by T cell polarization and flow cytometric analysis. **d,** Expression of CXCR5, FOXP3, and PD-1 in naïve CD4^+^ T cells cultured under Tfh-polarizing conditions (dark grey) or non-polarizing conditions (light grey). **e,** Log2 fold change relative to matched scramble controls for Th1 (left) and Th17 (right) cell frequency in TF-knockouts under Th1- or Th17-polarizing conditions. **f,** Log2 fold change relative to matched scramble controls for Tfh cell frequency in TF-knockouts under Tfh-polarizing conditions (gated on CD4^+^CD19^-^FOXP3^-^CXCR5^+^PD-1^+^ cells). Data were analyzed using a mixed effects model and Dunnett’s multiple comparisons test (**f**), one sample t test (**e**). Error bars SEM (**e-f**). \**P* < 0.05, \*\**P* < 0.01, \*\*\**P* < 0.001, \*\*\*\**P* < 0.0001.

To better understand this, we examined the differentially expressed genes (DEGs) between younger and older donors during early pseudotime stages. Early transcriptional programs in older individuals were enriched for stress- and effector-associated genes, including the gene encoding BLIMP-1 (*PRDM1*), which opposes BCL6 function, multiple AP-1 family transcription factors (*FOSB, JUNB, JUND, JUN*), and the AP-1 family binding partner *IRF4* (**Figure 6b**). In contrast, early transcriptional programs in younger individuals overexpressed TFs that promote CD4^+^ T helper cell plasticity and maintenance such as *BACH2, SOX4, LEF1,* and *STAT1*.

To functionally assess the roles of these age-associated TFs during early Tfh cell polarization, we developed an arrayed CRISPR-based polarization model using naïve CD4^+^ T cells. Following CRISPR/Cas9-mediated knockout (KO) of select age-associated TFs, naïve CD4^+^ T cells were cultured under Tfh-polarizing conditions for 7 days (**Figures 6c,d and S7a**). Transcription factors selected for knockout included canonical Tfh regulators (*BCL6, TCF7*) as well as age-associated transcriptional regulators identified from our trajectory analysis (**Table S4**). Samples demonstrating high editing efficiency along with a non-Tfh phenotype in response to *BCL6*-KO were retained for downstream analysis (**Figure S7b,c**). We additionally validated the CRISPR knockout platform by targeting key transcription factors required for alternative CD4^+^ T helper cell fates. Knockout of *STAT1* and *BATF* selectively disrupted Th1 and Th17 polarization, respectively, confirming that the assay faithfully captures lineage-specific differentiation (**Figures 6e and S7d,e**).

Among the transcription factors tested under Tfh-polarizing conditions, knockout of *BACH2* and *SOX4* significantly reduced Tfh cell differentiation, whereas *SP110* knockout produced a similar but not significant trend (*P*=0.07; **Figure 6f**). Knockout of *LEF1* or *TCF7*, the canonical regulators of the naïve-to-Tfh cell transition, showed effects in the expected direction but failed to achieve significance. Collectively, these results identify *BACH2* and *SOX4* as key regulators of early human Tfh cell polarization that are downregulated with age. These findings indicate that aging alters the transcriptional switch from activated precursors to committed Tfh cells, resulting in impaired progression to the effector CXCL13^+^ GC-Tfh cell state.

## Discussion

Durable antibody responses originate from germinal center B and T cell interactions, but our understanding of how these interactions are impacted by human aging has been incomplete. Here, we find that human Tfh cell development includes a mature CXCL13^+^ effector state that is absent in inbred mice. This distinct functional state features increased expression of the memory and tissue residency genes *IL6ST, IL7R, ITGAE,* and *TCF7* amongst others. Most importantly, we show that these specialized CXCL13^+^ GC-Tfh cells are strikingly reduced in older donors due to an early block in maturation. This loss of mature Tfh cells likely explains the poor antibody responses of older adults. By implicating defects in maturation rather than a generalized decline in B cell or Tfh cell frequencies, we provide a new framework for understanding why vaccines are less effective in older adults and highlight a human-specific mechanism of immune aging that could not be inferred from animal models alone.

A longstanding question has been whether impaired vaccine responses in older individuals arise primarily from intrinsic B cell dysfunction or from defective Tfh cell help. Studies in mice have highlighted defects in class-switch recombination and somatic hypermutation in aged B cells, but therapeutic interventions in humans suggest that B cells from older donors remain capable of responding when provided with adequate Tfh cell signals.^5,42,43^ Our findings support this, as peripheral B cells from older individuals responded to exogenous Tfh cell help signals similarly to B cells of younger donors. These findings position Tfh cells as the critical limiting component. Importantly, Tfh cells exist in phenotypically and functionally distinct states, yet the relative contribution of these states to GC function and their stability throughout the human lifespan has been unresolved. Among Tfh cell subsets, CXCL13⁺ GC-Tfh cells are emerging as particularly important regulators. CXCL13 is classically expressed by follicular dendritic cells and acts as a chemoattractant via the CXCR5 receptor to establish the canonical dark and light zone architecture. However, human Tfh cells can also express CXCL13, which murine T cells do not. In human studies, the abundance of CXCL13⁺ GC-Tfh cells correlates with vaccine responsiveness, and CXCL13 levels in blood following vaccination strongly correlates with GC activity and antibody quality.^17,18^ Given that the CXCL13^+^ GC-Tfh state is marked by significantly increased expression of the memory and tissue residency genes *TCF7, IL7R*, and *ITGAE* (encoding CD103), it is tempting to speculate that these cells are especially important drivers of long-term antibody-mediated immunity. Our study now positions these CXCL13⁺ GC-Tfh cells as a critical mature subset along the human Tfh differentiation continuum and shows that this maturation process is disrupted during aging. These results help clarify why previous reports of Tfh frequency in aging have been inconsistent, showing that the defect lies not in overall abundance but in the failure to efficiently produce mature GC-Tfh cells, especially those expressing CXCL13.

Furthermore, our study demonstrates that novel human biology can be discovered by combining thorough analyses of inter-human variability with *in vitro* models for experimental intervention. Although Tfh cells were previously identified as key contributors to the longitudinal human immune-aging metric IMM-AGE, peripheral blood is limited in its ability to serve as a relevant tissue to perform mechanistic studies.^16^ Here, by examining transcription factors downregulated with age, we uncovered novel regulators of human Tfh cell maturation. The transcriptional regulation of Tfh cell differentiation has been well characterized, largely by identification of regulators in mouse models followed by validation in human cells.^3,44–48^ These studies established the role of canonical Tfh regulators like BCL6, IRF4, and BATF. Our study extends this framework for human Tfh maturation beyond the initial stages of polarization and naïve CD4^+^ T cell commitment, revealing both conserved and divergent features. BACH2, for example, has been described in mice as a negative regulator that restrains Tfh fate,^49^ whereas in our human system it emerged as a positive regulator required for early progression toward the CXCL13⁺ GC-Tfh stage, underscoring stage- and species-specific differences. SOX4, previously implicated in driving CXCL13 expression in peripheral T cells during chronic inflammation,^50^ is shown to be necessary for GC-Tfh cell differentiation, linking a program associated with pathology to normal vaccine-induced humoral immunity. SP110, best characterized as a co-regulator of NF-κB signaling, was not previously associated with Tfh biology but in our study emerged as a potential regulator of human Tfh maturation. Notably, SP110 mutations are associated with primary immunodeficiency resulting in normal mature B cell frequencies, but lacking plasma and memory cells.^51,52^ These findings not only expand the transcriptional network that governs human Tfh cell development, but also highlight the importance of studying regulatory pathways. By combining *in vitro* models of human tonsils with CRISPR-based perturbations, we establish a mechanistic platform to dissect regulatory pathways directly in primary human model germinal centers. In this way, our system provides a framework for analysis of human immune responses to vaccination and offers a versatile tool to uncover human-specific checkpoints in adaptive immunity.

Our findings have translational potential for measuring the efficacy of vaccines during states of immunodeficiency. Plasma CXCL13 has already been characterized as a biomarker of GC activity in clinical vaccine studies. Future work will be necessary to determine the contributions of CXCL13^+^ GC-Tfh cells to systemic CXCL13 levels, and whether aging also directly impacts CXCL13^+^ Tfh cells in the periphery. Plasma CXCL13 may serve not only as a marker of GC integrity but also as a surrogate for the capacity of Tfh maturation, enabling stratification of older adults according to their likelihood of mounting effective vaccine responses in an accessible tissue compartment. The identification of transcriptional regulators such as BACH2 and SOX4 also points toward direct molecular targets for interventions designed to restore Tfh cell maturation, whether through adjuvant formulations, cytokine manipulation, or pharmacological alterations of transcriptional networks.^15,53^ At the same time, several limitations should be noted. Tonsil tissue is an accessible source of mucosal secondary lymphoid B and T cells, however they may not fully reflect the biology of lymph nodes or spleen. Additionally, our organoid system recapitulates many features of GC function, though it cannot capture systemic trafficking or long-term recall effects that happen *in vivo*. Although single TF perturbations provide important insights, Tfh lineage commitment likely emerges from the interplay of multiple transcription factors. Our approach therefore captures only part of the picture, as combinatorial knockouts of multiple TFs could either exacerbate or counterbalance the effects we observed, revealing new layers of regulation yet to be uncovered. Lastly, our CRISPR screen was focused rather than unbiased, and additional regulators are likely to contribute to Tfh maturation *in vivo*. Despite these caveats, the integration of human immune organoids, single-cell profiling, and genetic perturbations provides a powerful framework for uncovering mechanisms of adaptive immunity that are inaccessible in animal models.

In conclusion, our study identifies a human-specific population of mature CXCL13-producing GC-Tfh cells that is diminished by an aging-associated block in Tfh maturation, reshaping our understanding of the mechanisms underlying impaired antibody responses in older adults. By highlighting the deficit in early Tfh cell maturation and identifying molecular regulators, we establish a foundation for future strategies aimed at restoring robust immunity in aging populations.

## Methods

### Sample collection

Whole tonsils from 55 consented patients undergoing clinically indicated surgery for obstructive tonsillar hypertrophy (sleep disordered breathing, excluding cases of recurrent tonsillitis) were collected at Stanford Hospital, Palo Alto, USA and Alberta Children’s Hospital, Calgary, Canada (**Table S1**). Ethics approval was granted by the Stanford University Institutional Review Board (IRB60741 and IRB30837) and University of Calgary Research Ethics Board (REB24-1330). Written informed consent was obtained from adult participants or from the legal guardians for pediatric patients. Medications were reviewed to ensure there was no previous use of immunosuppressive therapies. In our cohort, 21 participants were aged 5-17 years (male n = 8, female n = 13) and 34 were adults of 19-73 years (male n = 23, female n = 11).

Extracted whole tonsils were collected immediately after surgery in saline solution and kept on ice, then rinsed in PBS (Gibco) and incubated > 30 min at 4°C in cold decontamination medium: Ham’s F-12 (Gibco) containing 100 µg/mL Normocin (InvivoGen), and 200 U/mL Penicillin-Streptomycin (Pen-Strep; Gibco). After decontamination, tonsils were rinsed with PBS and processed as described below.^21^

### Human vaccination protocol and hemagglutination inhibition assay

Serological data were obtained from healthy volunteers enrolled across several related influenza vaccination studies conducted through the Stanford-Lucille Packard Children’s Hospital (LCPH) vaccine research program as previously described.^24^ All protocols were approved by the Stanford University Institutional Review Board. Written informed consent was obtained from each participant. Participants were generally healthy according to self-reporting, with no acute or chronic medical conditions. Individuals with any known or suspected impairment in immune function were excluded, including those with history of immunodeficiency, clinically significant liver disease, insulin-treated diabetes, moderate to severe kidney disease, chronic hepatitis B or C virus, or blood pressure greater than 150/95 mmHg at screening. Subjects were also excluded if they received blood products within 6 months or donated blood within 6 weeks, or showed any features of viral infection at baseline sampling. None of the participants had exposures to immunosuppressive medications.

Peripheral blood was collected at day 0 before vaccination, day 7 (+/-1), and day 28 (+/-4) after a single intramuscular dose of the inactivated seasonal influenza vaccine. Serum antibody titers against H1, H3, and B influenza antigens were measured using a standard hemagglutination inhibition assay. For each donor, geometric mean titers across all antigens were calculated, and fold-changes from baseline were used to assess vaccine induced responses. Seroconversion was defined as ≥ 4-fold increase in HAI with day 28 GMT ≥ 40. Seroprotection was defined as day 28 GMT ≥ 40. Associations between age and vaccine response were modeled using linear and logistic regression, controlling for baseline titer and sex. The cohort included 402 participants (aged 8-90 years; female/male ratio 1.5:1). Additional cohort information and data access are available at DOI: 10.1038/s41597-019-0213-4.

### Human tonsil tissue and blood processing

Tonsil tissue was mechanically dissociated and passed through a 100-μm cell strainer using a syringe plunger into complete medium: RPMI with GlutaMAX (Gibco), 10% fetal bovine serum (FBS; R&D Systems), 100 U/mL Pen-Strep, 100 µg/mL Normocin, MEM non-essential amino acids (MEM-NEAA; Gibco), 1 mM sodium pyruvate (Gibco), and Insulin-Transferrin-Selenium (Gibco). The cell suspension was purified by Ficoll density gradient separation (Cytiva) and washed in complete medium. Isolated cells were frozen in FBS containing 10% DMSO (Sigma).

Peripheral blood of 27 healthy blood donors aged 16-72 years (male n = 20, female n = 7) were provided by the Stanford Blood Center. Informed consent was obtained in accordance with the Declaration of Helsinki protocol. Donor characteristics are provided in **Table S1**. PBMCs were isolated by density gradient centrifugation and cells were frozen as described above. All cryopreserved cells were maintained in liquid Nitrogen until further use.

### Cell culture and organoid preparation

Cryopreserved tonsil cell aliquots were thawed in a 37°C water bath into complete medium, enumerated, and resuspended to 6 × 10^7^ cells/mL. Cells were embedded in a hydrogel matrix (VitroGel ORGANOID-1; The Well Bioscience) to improve throughput and increase technical replicates. Organoids were assembled to 2.5 x 10^6^ cells in 15 µL droplets and plated under sterile conditions into 96-well V-bottom plates containing 150 µL prewarmed complete medium supplemented with 0.5 μg/mL of recombinant human B cell-activating factor (BAFF; Biolegend) to improve B cell survival and overall cell recovery in the organoid. Fresh culture media was replenished on day 4, 7, and 10. For antigen stimulation, 0.5 µL PBS or LAIV (containing 0.16-1.6 x 10^5^ fluorescent focus units per strain; FLUMIST QUADRIVALENT, MedImmune LLC) was added to each well.

### ELISA for detection of HA-specific antibodies

Clear flat-bottom polystyrene high-binding 96-well microplates (Corning) were coated with 100 ng/well of recombinant influenza A H1N1 hemagglutinin protein (A/California/7/2009; Sino Biological) in 100 mM bicarbonate buffer (pH 9.6) overnight at 4°C. All washing steps were done on a BioTek 450TS microplate washer (Agilent). Coated plates were washed 3x with PBS containing 0.05% Tween20 (PBS-T; Sigma) and blocked with PBS containing 1% BSA (Miltenyi) for 1 h at room temperature (RT). Culture supernatants were added undiluted or as 1:10, 1:100, and 1:300 dilution in PBS and incubated for 1 h at RT. Following 3x washing steps with PBS-T, plates were incubated with 1 ng/well of horseradish peroxidase-conjugated F(ab’)2 goat anti-human IgG-Fc cross-adsorbed secondary antibody (Bethyl) for 1 h at 37°C. For detection, plates were washed 3x with PBS-T and 1-step Ultra TMB substrate solution (Thermo Fisher Scientific) was added for ∼10 min at RT. After quenching with sulfuric acid, absorbance at 450 nm was recorded on a BioTek Cytation 7 multimode reader (Agilent). Absorbance values were normalized for variations in cell counts to ensure accurate comparisons across samples. Data was analyzed using the ‘flux’ package (v.0.3.0) to visualize and calculate the area under the curve for each condition’s dilution series.

### Biolayer Interferometry

Organoids were grown in low (1 x 10^6^) and high (5 x 10^6^) cell density as triplicates in 12-well plates in the presence or absence of LAIV for 14 days. Supernatants were collected on day 4, 7, 10, and 14 and pooled across technical replicates before readout. The supernatants were then processed using 3x exchanges in PBS with 50 kDa cut-off spin columns (Amicon) and concentrated to a final volume of 200 µL each. Binding affinities were calculated as previously described using an Octet Qk instrument (Pall ForteBio).^21^ Briefly, full-length H1 CA/09 HA antigen was expressed and purified using Ni-NTA affinity columns followed by size-exclusion cleanup.^54,55^ Purified antigens were captured on AR2G biosensor tips using the amine reactive second-generation reagent kit. The ligand-bound biosensors were dipped into 2-fold serially diluted, processed antibody supernatants. The association and dissociation were both monitored for 1 h. PBS-T (pH 7.4) was used for sample dilution. Dissociation was carried out in PBS. Double referencing was performed using unligated biosensors and an irrelevant E.coli maltose-binding protein. The dissociation rate constant K_off_ per minute was determined by a global fit of exponential decay kinetics. Each binding interaction was performed in duplicates.

### B-Tfh cell co-culture

Co-cultures of Tfh and B cells were prepared by magnetic-activated cell sorting (MACS) followed by fluorescent-activated cell sorting (FACS) from cryopreserved tonsil single-cell suspensions. In brief, cells were thawed in a 37°C water bath and CD3^+^ T cells were enriched using the human Pan T Cell Isolation Kit (Miltenyi). The retained fraction enriched to contain ∼90% B cells was collected and kept at 37°C in complete medium. The enriched CD3^+^ fraction was incubated with Human TruStain FcX (Biolegend) for 10 min on ice and stained in FACS buffer (PBS, 2 mM EDTA, 0.5% BSA or 2% FBS) containing anti-human CD4, CD8, CD14, CD19, CD25, CXCR5, and PD-1 antibodies for 30 min on ice. Samples were used to determine the ratio of Tfh cells (CD4^+^CD8^-^CD19^-^CD14^-^CD25^-^CXCR5^+^PD-1^+^) to Tfr (CD4^+^CD8^-^CD19^-^CD14^-^CD25^+^CXCR5^mid^PD-1^mid^). Tfh cells were sorted on a FACS Aria Fusion cytometer (BD Biosciences) directly into sterile complete media following the gating strategy in **Figure S1b**. Sorted cells were washed in complete media, counted, and diluted. Enriched B cells (5 x 10^4^) and Tfh cells (3 x 10^4^) were plated in U-bottom 96-well plates containing 150 µL complete media supplemented with BAFF and stimulated with 0.5 µL/well of LAIV. Plasmablasts (CD3^-^/CD38^hi^/CD27^hi^) were analyzed on day 7 by flow cytometry (**Figure S1c**) on a NovoCyte Penteon cytometer (Agilent).

### B cell activation *in vitro*

Cryopreserved PBMCs were thawed in a 37°C water bath into complete medium and B cells were enriched using the EasySep Human B Cell Isolation Kit (StemCell) according to the manufacturer’s instructions and plated in 96-well flat-bottom plates at 2 x 10^5^ cells in 200 µL complete medium containing 100 U/mL IL-2 (R&D). Cells were stimulated with 6 µg/mL of the BCR agonist (F(ab’)₂ Anti-IgA/IgG/IgM; Jackson ImmunoResearch) with or without the T cell help signals IL-21 (50 ng/mL; proteintech) and MEGACD40L (100 ng/mL; Fisher Scientific). IL-2 and IL-21 were supplemented on day 2 and day 4 by exchanging 50 µL of medium per well. Stimulation protocol was adapted from Cocco et al.^56^ Cells were harvested after 7 days and stained for flow cytometry with LIVE/DEAD Fixable Blue (Thermo Fisher) and anti-human CD19, CD27, CD38, CD138, CD95, and IgD antibodies in the presence of Human TruStain FcX. Stained samples were analyzed on a NovoCyte Penteon cytometer (**Figure S2**).

### Single-cell RNA-sequencing

#### Library preparation, sequencing, and alignment

Cryopreserved tonsil single-cell suspensions for each donor were thawed in a 37°C water bath and magnetically sorted using the Pan T Cell Isolation Kit (Miltenyi). For each sample, 2 x 10^6^ enriched CD3^+^ T cells were resuspended in FACS buffer with Human TruStain FcX for 10 min and stained for 30 min with TotalSeqC antibodies against CD3, CD4, and CD8 (Biolegend). Each sample was washed twice with FACS buffer and resuspended in PBS containing 0.04% BSA. Tonsil samples for each donor were counted and diluted for scRNA-seq cell capture on a 10x Genomics Chromium Controller targeting 10,000 cells. Individual gene expression (GEX) and Antibody-Dependent Tag (ADT) libraries were prepared according to the Chromium Next GEM Single Cell 5’ Reagents Kit v2 protocol (10x Genomics). Libraries were sequenced on an Illumina Novaseq6000 using 151/10/10/151 base pair configuration (Novogene). Demultiplexed FASTQ files were aligned to the hg38 reference genome (GRCh38-2020-A) and processed using the ‘cellranger multi’ command.

#### Quality control, integration, and cell annotation for tonsil scRNA-seq datasets

Analysis was done in R version 4.2.2 using ‘Seurat’ (v.4.3.0). To demultiplex tonsil samples for each donor, hashtag oligo (HTO) counts were first normalized with centered log-ratio transformation (CLR) and separated with HTODemux (positive.quantile = 0.99). Tonsil samples from each donor were then pre-processed independently to keep cells containing 200-6,000 genes with < 10% of counts containing mitochondrial genes. Doublets were identified and removed using ‘DoubletFinder’ v.2.0.3.^57^

Donor-specific genes were first removed from the matrices, including TRAV, TRBV, and HLA. Datasets were then normalized using ‘SCTransform’, cell cycle effects were regressed, and the top 3,000 highly variable genes were used for downstream dimensionality reduction with Principal Components Analysis (PCA). Jackstraw plots were used to determine the number of principal components for further analysis (dims = 1:20). These dimensions were used to construct a 2-dimensional representation using uniform manifold approximation and projection (UMAP, n.neighbors = 20, min.dist = 0.35). We identified minor batch effects attributed to individual donors (**Figure S3c,d**) and therefore integrated the datasets using ‘Harmony’ (v.1.2.0, group.by.vars = “sample”).

The top 20 harmony dimensions were used to identify clusters (resolution = 0.8). For cell classification, cluster marker genes were reviewed with Seurat’s ‘FindAllMarkers’ and each population was assessed to ensure representation for multiple donors.

### Flow cytometric analysis of tonsil Tfh cells

Cryopreserved tonsil cells were thawed and stained with LIVE/DEAD Fixable Blue and anti-human CD19, CD3, CD4, CXCR5, PD-1, and ICOS antibodies; followed by fixation and permeabilization with FOXP3/Transcription Factor Staining Buffer Set (Thermo Fisher) and intracellular staining for FOXP3. Cells from the same donors were stimulated overnight with PMA/ionomycin (Thermo Fisher) to measure CXCL13 production. For ICS, cells were first stained with LIVE/DEAD Fixable Blue and anti-human CD45, CD14, CD16, CD19, TCRαβ, CD8, CD4, CXCR5, PD-1, and CD25 antibodies; followed by Cytofix/Cytoperm Fixation/Permeabilization Solution Kit (BD) and intracellular staining for CXCL13. All stains were performed with Human TruStain FcX. Samples of < 45% viability were excluded. Stained samples were analyzed on a NovoCyte Penteon cytometer (**Figure S6a**).

### ELISA for quantification of CXCL13 and IL-21 in cell culture supernatants

CXCL13 was quantified using the BLC/CXCL13 human ELISA Kit or the IL-21 human ELISA kit (Invitrogen) according to the manufacturer’s instructions. Cell culture supernatants were diluted 1:10 in the respective diluent. Concentration was determined using standards (0-1000 pg/mL). Absorbance at 450 nm was recorded on a BioTek Cytation 7 multimode reader (Agilent). Values were normalized for variations in cell counts to ensure accurate comparisons across samples.

### Tfh cell trajectory analysis in old vs. young donor samples

We mapped a trajectory within sub-clustered CD4⁺ T cell populations using slingshot (v2.6.0). We specified a biologically informed trajectory starting from naïve CD4⁺ T cells and ending in germinal center T follicular helper cells, with intermediate states including activated CD4^+^ T cells and Tfh cells. Pseudotime values along each inferred lineage were extracted using ‘slingPseudotime’ for visualization and downstream analysis. We used tradeSeq (v1.14.0) to identify genes whose expression dynamics varied significantly along the CD4⁺ T naïve - CD4⁺ T activated - CD4⁺ Tfh - CD4⁺ GC-Tfh trajectory. Gene-level count data were extracted from the RNA assay, restricted to the set of highly variable genes defined in the SCT assay.

### Analysis of differentially expressed genes along pseudotime

To analyze DEGs across pseudotime, we implemented a sliding-window differential expression approach. In brief, individual cells were ordered by pseudotime, and overlapping windows of fixed size (width = 1 pseudotime unit, step = 0.1 pseudotime unit) were defined across pseudotime. Within each window, we compared younger vs. older samples using the Wilcoxon rank-sum test in Seurat’s “FindMarkers” function. DEGs were defined as those with an adjusted *P* ≤ 0.05. The number of DEGs was recorded for each window and plotted across pseudotime, which enabled identification of regions along the trajectory with a high density of DEGs. Based on these profiles, two pseudotime regions enriched in DEGs were defined as “early” and “late”.

### Knockout of transcription factors using CRISPR/Cas9

Cryopreserved tonsil cells were thawed in a 37°C water bath into serum-free medium containing 10 µg/mL DNAse (Roche) and incubated for ∼1 h at 37°C and 5% CO_2_. Knockout was performed using chemically modified synthetic single guide RNAs (gRNAs) formulated in proprietary gene knockout kits (Synthego/EditCo; **Table S2**) targeting genes encoding the selected transcription factors. Knockout of *BCL6* served as a positive control and scramble gRNA #1 was used as a negative control. Ribonucleoproteins (RNPs) were formed by gently pipetting Cas9 protein (Aldevron) to gRNA in a molar ratio of 1:2 (Cas9:gRNA). Following incubation at RT for 15 min, RNPs were kept on ice until use. After resting, tonsil cells were washed in PBS (400x g, 5 min, RT) and resuspended to 4-15 x 10^7^ cells/mL in P3 Primary Cell Nucleofection Solution (Lonza). For electroporation, 20 µL cells were mixed with 2.2 µL of each RNP (60 pmol Cas9 + 120 pmol gRNA) and transferred to one well of a Nucleocuvette Strip (4D-Nucleofector X kit S; Lonza). Cells were electroporated using the EH-100 pulse program on a 4D-Nucleofector (Lonza), rescued by immediately adding 80 µL/well of pre-warmed serum-free medium, and incubated at 37°C and 5% CO_2_ for ∼30 min. Following the resting period, cells were transferred to 96-well U-bottom plates and Nucleocuvette wells were rinsed with 100 µL complete medium containing 20% FBS and 1 µg/mL BAFF. Each well was supplemented with 100 µL/well of complete medium with LAIV (0.5 µL/well). On day 2, 3, and 5, wells were replenished with 50 µL/well complete medium with BAFF.

On day 7, supernatants for ELISA were collected, snap-frozen, and kept at -80°C until use. Cells were harvested and stained with LIVE/DEAD Fixable Blue Cell Stain and anti-human CD3, CD4, CD19, CD25, CD27, CD38, CXCR5, and PD-1 antibodies as described previously and analyzed on a NovoCyte Penteon cytometer.^58^

### Naïve CD4^+^ T cell polarization and transcription factor knockout

Cryopreserved PBMCs were thawed in a 37°C water bath into complete medium and naïve CD4^+^ T cells were enriched using the EasySep Human naïve CD4^+^ T Cell Isolation Kit (StemCell) according to the manufacturer’s instructions. CRISPR/Cas9-based knockout was performed as described above. After ∼1 h of rescue at 37°C, edited naïve CD4^+^ T cells were seeded in 96-well flat-bottom plates to 40,000 cells/well in complete media and polarized by adding ImmunoCult only (Th0; StemCell) or ImmunoCult with the following cytokines and neutralizing antibodies: Th1 polarization cocktail (StemCell) for Th1 polarization; IL-1b (f.c. 10 ng/mL), IL-6 (f.c. 10 ng/mL), IL-21 (f.c. 50 ng/mL), IL-23 (f.c. 20 ng/mL), TGFß1 (f.c. 5 ng/mL), anti-IL-4, and anti-IFN𝛾 (f.c. 10 µg/mL, all from proteintech) for Th17 polarization. For Tfh cell polarization, a 96-well flat-bottom plate was coated overnight at RT with 100 µL/well of anti-CD40L recombinant monoclonal antibody (clone hu5c8, f.c. 5 µg/mL; Enzo) and recombinant human B7-H2 Fc chimera protein (f.c. 5 µg/mL, R&D) and seeded naïve CD4^+^ T cells were supplemented with ImmunoCult, IL-6 (f.c. 100 ng/mL), IL-21 (f.c. 50 ng/mL), anti-IL-2 (f.c. 5 µg/mL, R&D), and anti-IFN𝛾 (f.c. 10 µg/mL). Cells were cultured for 7 days and complete medium without cytokines or antibodies was replenished on day 2, 4, and 6. On day 7, cells were harvested and stained with LIVE/DEAD Fixable Blue Stain and anti-human CD4, CCR4, CCR6, CXCR3, CXCR5, and PD-1 as well as TBET and FOXP3 antibodies for Th0-, Th1-, and Tfh-polarized cells or GATA3 and FOXP3 for Th0- and Th17-polarized cells (**Figure S7**). All cells were analyzed on a NovoCyte Penteon cytometer.

### Assessing CRISPR editing efficiency

Genomic DNA was extracted using the QuickExtract solution (LGC Biosearch Technologies). To amplify the genetic region of interest, polymerase chain reaction (PCR) was performed using gene-specific oligonucleotides (ELIM Biopharm) with the Platinum II Hot-Start PCR Master Mix (Invitrogen) according to the manufacturer’s instructions on a Mastercycler pro (Eppendorf). Successful PCR amplification was confirmed by agarose gel electrophoresis and PCR amplicons were transferred to ELIM Biopharm for Sanger sequencing. Editing efficiencies (**Table S3**) were calculated using the Inference of CRISPR Edits (ICE) analysis tool (Synthego Performance Analysis 2019, v3.0) to ensure target gene knockout. Oligonucleotides used for PCRs and sequencing are listed in **Table S2**.

### Software and statistical analysis

Unless otherwise stated, all analyses were performed using R version 4.2.2. Individual statistical tests for various analyses are listed in their respective figure legends. There were no formal statistical tests to predetermine sample sizes, and investigators were not blinded to sample conditions during the analysis. For CRISPR knockout experiments, one donor was excluded from all conditions due to consistently reproducible poor knockout efficiencies. FlowJo v.10.10.0 was used to analyze flow cytometry data. Synthego Performance Analysis, ICE Analysis tool. 2019. v.3.0 was used to assess CRISPR editing efficiencies. GraphPad Prism 10 was used for statistical analysis of data from flow cytometry experiments and CRISPR knockouts. BioRender was used to create illustrations for all schematics.

## Resource Availability

### Data availability

Single-cell RNA-seq data have been deposited at GEO (GSE270554) and are publicly available as of the date of publication. Any additional information is available from the lead contact Mark M. Davis upon request.

### Materials availability

This study did not generate new unique reagents.

## Acknowledgments

We thank Lei Chen for processing tonsil and peripheral blood samples and all members of the Davis lab for critical input and discussion. Cell sorting was done in the Stanford Shared FACS Facility on the FACSAria Fusion purchased by the Parker Institute for Cancer Immunotherapy. We thank the German Research Foundation (DFG) for funding the Walter Benjamin Program (T.B.). This research was supported by grants from the Open Philanthropy Foundation, the National Institute of Health (NIH 3U19AI057229-17S1, NIH P01 AI153559-01, 1F32AI186256-01 (C.B.)), Israel Science Foundation (grant #839/21), and the Howard Hughes Medical Institute.

## Author contributions

Conceptualization, N.A.B., C.B., T.B., and M.M.D.; Methodology, N.A.B., C.B., T.B., A.B., T.J.F.-C., and S.S.S.-O.; Software, N.A.B., A.H.L., A.M., and V.S.; Validation, N.A.B., C.B., and T.B.; Formal Analysis, N.A.B., C.B., T.B., A.B., and T.J.F.-C.; Investigation, N.A.B., C.B., and T.B.; Resources, E.S., L.K.; Data Curation, N.A.B., A.H.L., A.B., T.J.F.-C., A.M., V.S., V.M., R.C., and S.S.S.-O.; Writing – Original Draft, N.A.B., C.B., and T.B.; Writing – Review & Editing, N.A.B., C.B., T.B., A.H.L., E.S., A.B., T.J.F.-C., A.M., V.S., V.M., L.K., R.C., S.S.S.-O. and M.M.D.; Visualization, N.A.B., C.B., T.B., and A.B.; Supervision, S.S.O.-O and M.M.D.; Funding Acquisition, S.S.S.-O. and M.M.D..

## Supplemental items

Figure S1. Tfh:Tfr ratio and flow cytometry gating strategies. Related to **Fig. 1**.

Figure S2. Flow cytometry gating strategy for B cell subsets. Related to **Fig. 2**.

Figure S3. scRNA-seq quality control and CXCL13^+^ GC-Tfh cells. Related to **Fig. 3**.

Figure S4. Validation in human tonsil atlas dataset.^36^ Related to **Fig. 3**.

Figure S5. Comparison of CD4^+^ T cell gene signatures. Related to **Fig. 4**.

Figure S6. Flow cytometry gating strategies and supporting data. Related to **Fig. 5**. Figure S7. Flow cytometry gating strategies and quality control. Related to **Fig. 6**.

Table S1. Donor characteristics. Related to all Figures.

Table S2. Oligonucleotide and gRNA sequences. Related to **Figs. 4i and 6e,f**.

Table S3. CRISPR editing efficiencies determined by ICE. Related to **Figs. 4i & 6e,f**.

Table S4. List of differentially expressed genes. Related to **Fig. 6f**.

