## Supplementary Figures for "Aging restricts maturation of CXCL13^+^ T follicular helper cells in human immunity"

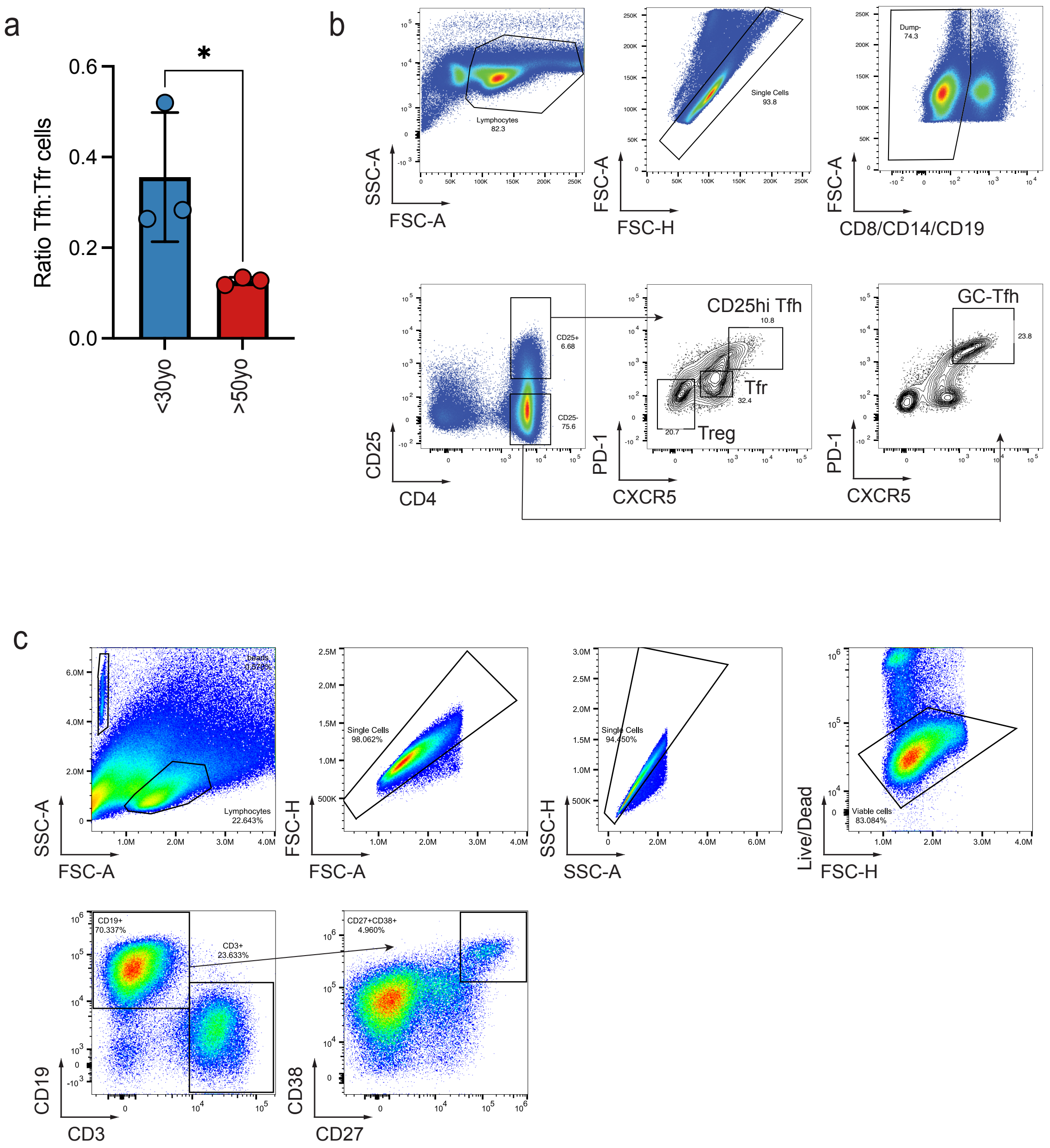


**Figure S1. Tfh:Tfr ratio and flow cytometry gating strategies. Related to Fig. 1.**

**a**, Tfh:Tfr ratios of three younger (blue, left) and three older (red, right) donors.

**b,** Gating strategy for sorting GC-Tfh cells ($CD4^{+}CD8^{-}CD14^{-}CD19^{-}CD25^{-}CXCR5^{hi}PD\text{-}1^{hi}$) and Tfr cells ($CD4^{+}CD8^{-}CD14^{-}CD19^{-}CD25^{+}CXCR5^{mid}PD\text{-}1^{mid}$).

**c**, Gating strategy for assessment of plasmablast frequencies ($CD3^{-}CD19^{+}CD27^{hi}CD38^{hi}$).

Data in **a** were analyzed using one-way ANOVA with post-hoc Tukey test. *$P < 0.05$.

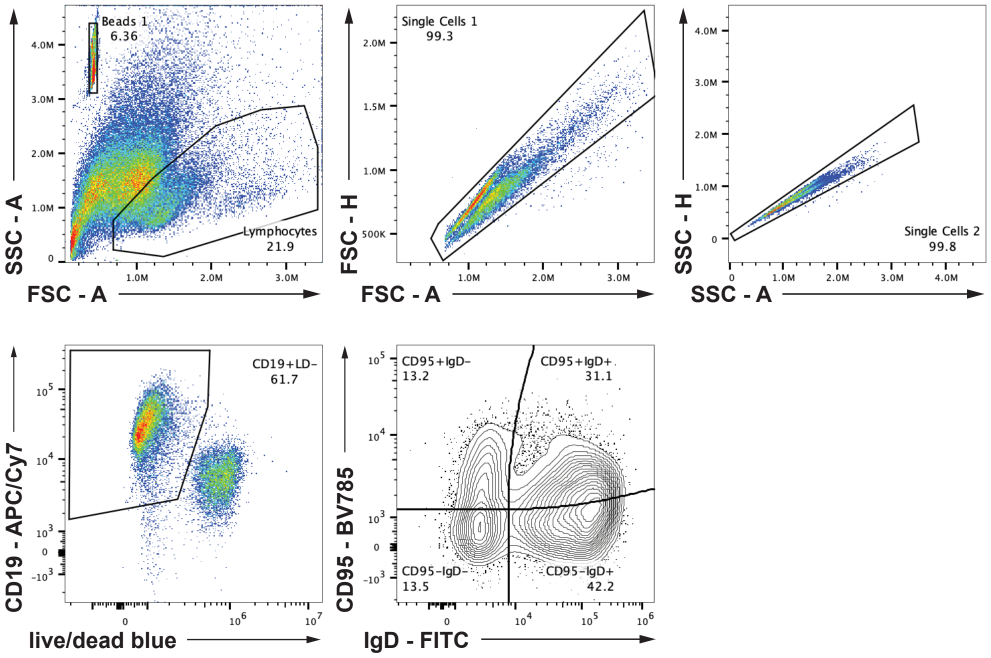


**Figure S2. Flow cytometry gating strategy for B cell subsets. Related to Fig. 2.**

Exemplary gating strategy for different B cell subsets.

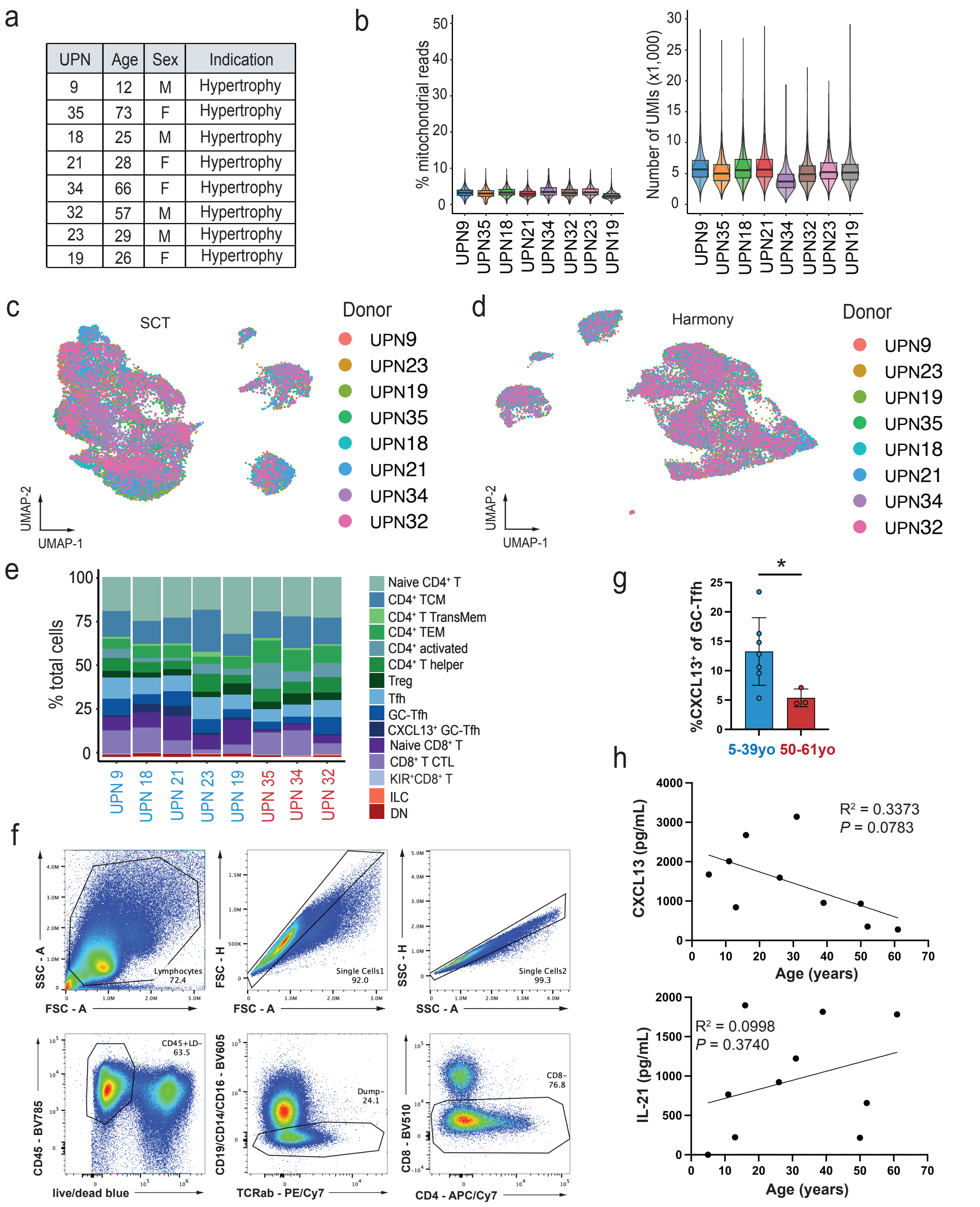


**Figure S3. scRNA-seq quality control and CXCL13+ GC-Tfh cells. Related to Fig. 3.**

**a,** Donor characteristics (unidentified patient numbers (UPNs), age in years, sex, and indication for tonsillectomy) for samples used for scRNA-seq analysis.

**b,** Mitochondrial gene reads in scRNA-seq from tonsils (left) and Unique Molecular Identifiers (UMIs, right) in scRNA-seq data from tonsil samples (n = 8).

**c,** Uniform Manifold Approximation and Projection (UMAP) of PCA reduction demonstrating donor-level batch effects in scRNA-seq.

**d,** UMAP representation of harmony reduction demonstrating improvement in batch effects.

**e,** Compositional frequencies for young (blue) and old (red) tonsil T cell populations.

**f,** Gating strategy for tonsil $CD4^+$ cells ($CD45^+CD14^-CD16^-CD19^-CD8^-$).

**g,** Percentage of $CXCL13^+$ GC-Tfh cells in 5-39yo (blue, n = 7) vs. 50-61yo donors (red, n = 3).

**h,** Linear regression analysis examining the correlation between age and CXCL13 (top) or IL-21 secretion (bottom; n = 10).

Data in **g** were analyzed using Mann-Whitney test. **P* < 0.05.

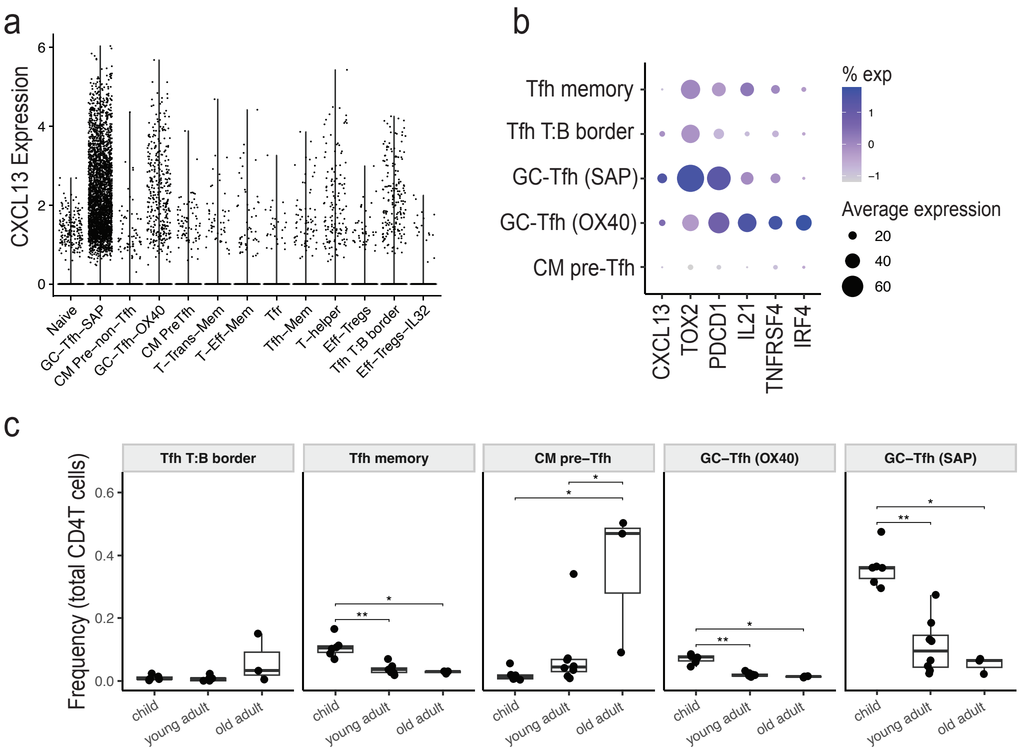


**Figure S4. Validation in human tonsil atlas dataset.[36] Related to Fig. 3.**

**a,** CXCL13 gene expression across $CD4^+$ T cell populations from the human tonsil atlas.

**b,** Dotplot showing expression of selected genes across Tfh cell subsets from the human tonsil atlas. Dot size represents the fraction of cells expressing each gene, color indicates scaled average expression. GC-Tfh (SAP) cells are enriched for the CXCL13-effector program, whereas GC-Tfh (OX40) cells preferentially express an IL-21-associated activation program.

**c,** Donor-level frequency of Tfh cell subsets across age groups from the human tonsil atlas. Statistical comparisons show pairwise Wilcoxon rank-sum tests with Benjamini-Hochberg correction, *P < 0.05, **P < 0.005.

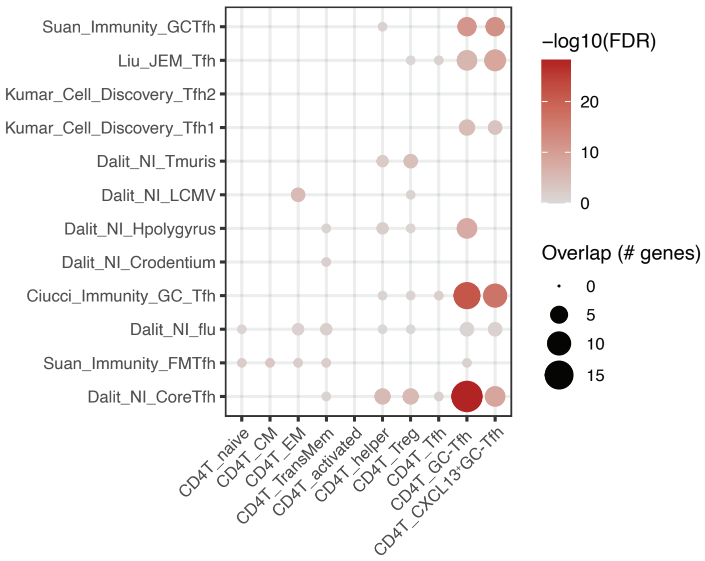


**Figure S5. Comparison of CD4$^+$ T cell gene signatures. Related to Fig. 4.**

Comparison of published murine Tfh cell gene signatures with CD4$^+$ T cell profiles from the human aging T cell scRNA-seq dataset.

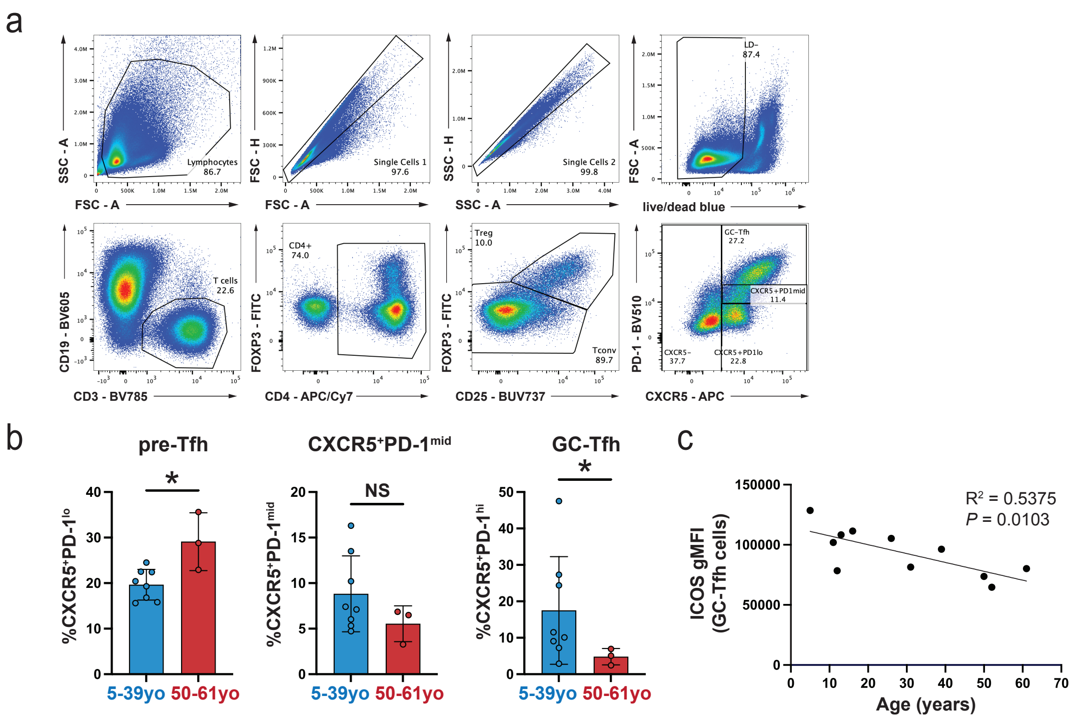


**Figure S6. Flow cytometry gating strategy and supporting data. Related to Fig. 5.**

**a,** Gating strategy for Tfh cell subsets ($CD3^+CD19^-CD4^+FOXP3^-CD25^-$) along their activation-to-differentiation trajectory.

**b,** Percentage of Tfh cell subsets in 5-39yo (blue, n = 8) vs. 50-61yo donors (red, n = 3) for $CXCR5^+PD\text{-}1^{lo}$ pre-Tfh cells (left), $CXCR5^+PD\text{-}1^{mid}$ Tfh cells (middle), and $CXCR5^{hi}PD\text{-}1^{hi}$ mature GC-Tfh cells (right).

**c,** Linear regression analysis examining the relationship between age and geometric mean fluorescence intensity of ICOS (n = 11).

Data in **b** were analyzed using Mann-Whitney test. *P < 0.05, **P < 0.01, NS, not significant.

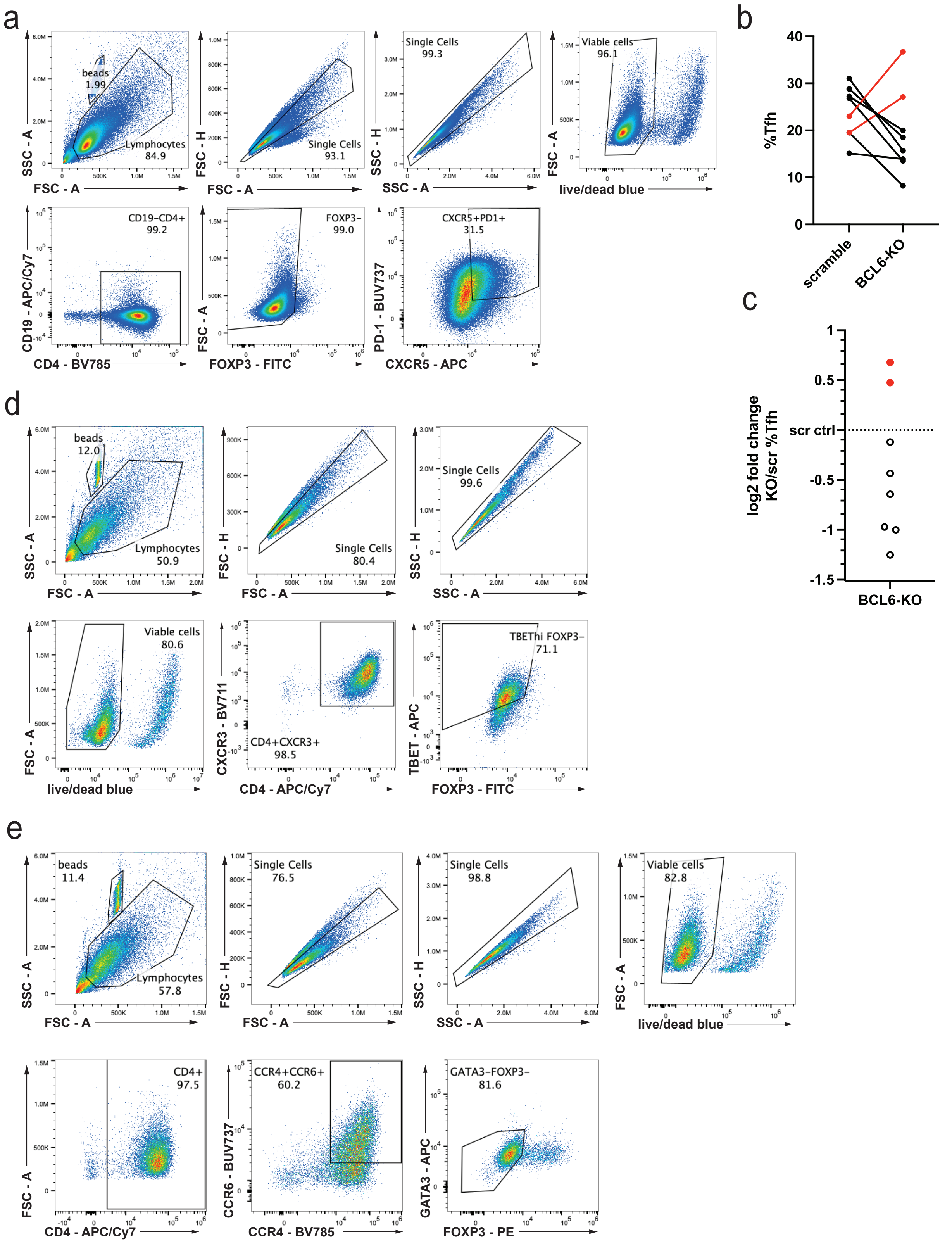


**Figure S7. Flow cytometry gating strategies and quality control. Related to Fig. 6.**

**a,** Gating strategy for Tfh cells induced from naïve CD4$^+$ T cells (CD4$^+$CD19$^-$FOXP3$^-$CXCR5$^+$PD-1$^+$).

**b,** Quality control (QC failed = red) for Tfh cell frequencies in BCL6-KO vs. scramble control under Tfh-polarizing conditions ($CD4^+CD19^-FOXP3^-CXCR5^+PD\text{-}1^+$).

**c,** Log2 fold change relative to matched scramble controls for Tfh cell frequency in BCL6-KO under Tfh-polarizing conditions ($CD4^+CD19^-FOXP3^-CXCR5^+PD\text{-}1^+$). Donors failed QC (red) when log2 fold change value was > 0.

**d,** Gating strategy for Th1 cells ($CD4^+CCR3^+FOXP3^-$ $TBET^{hi}$).

**e,** Gating strategy for Th17 cells ($CD4^+CCR4^+CCR6^+FOXP3^-GATA3^-$).
